# Functional 3D architecture in an intrinsically disordered E3 ligase domain facilitates ubiquitin transfer

**DOI:** 10.1101/831362

**Authors:** Paul Murphy, Yingqi Xu, Sarah L. Rouse, Steve J. Matthews, J Carlos Penedo, Ronald T. Hay

## Abstract

Post-translational modification of proteins with ubiquitin represents a widely used mechanism for cellular regulation. Ubiquitin is activated by an E1 enzyme, transferred to an E2 conjugating enzyme and covalently linked to substrates by one of an estimated 600 E3 ligases (1). RING E3 ligases play a pivotal role in selecting substrates and priming the ubiquitin loaded E2 (E2~Ub) for catalysis (2,3). RING E3 RNF4 is a SUMO targeted ubiquitin ligase (4) with important roles in arsenic therapy for cancer (4,5) and in DNA damage responses (6,7). RNF4 has a RING domain and a substrate recognition domain containing multiple SUMO Interaction Motifs (SIMs) embedded in a region thought to be intrinsically disordered (8). While molecular details of SUMO recognition by the SIMs (8–10) and RING engagement of ubiquitin loaded E2 (3,11–15) have been determined, the mechanism by which SUMO substrate is delivered to the RING to facilitate ubiquitin transfer is an important question to be answered. Here, we show that the intrinsically disordered substrate-recognition domain of RNF4 maintains the SIMs in a compact global architecture that facilitates SUMO binding, while a highly-basic region positions substrate for nucleophilic attack on RING-bound ubiquitin loaded E2. Contrary to our expectation that the substrate recognition domain of RNF4 was completely disordered, distance measurements using single molecule Fluorescence Resonance Energy Transfer (smFRET) and NMR paramagnetic relaxation enhancement (PRE) revealed that it adopts a defined conformation primed for SUMO interaction. Mutational and biochemical analysis indicated that electrostatic interactions involving the highly basic region linking the substrate recognition and RING domains juxtaposed those regions and mediated substrate ubiquitination. Our results offer insight into a key step in substrate ubiquitination by a member of the largest ubiquitin ligase subtype and reveal how a defined architecture within a disordered region contributes to E3 ligase function.

## Main

Previous NMR analysis of RNF4 revealed an N-terminal domain containing the 4 SIMs with poor chemical shift dispersion, indicative of a disordered region (8). Analysis of a sub-region encompassing SIMs 2+3 bound to diSUMO showed that the SIM 2+3 region adopts a “kinked” conformation that restrains the two linked SUMOs (8). We hypothesised that in the absence of SUMO chains the SIM containing region exhibits an extended, disordered conformation that becomes more ordered and compact upon binding to SUMO chains. To test this hypothesis, we performed smFRET measurements to determine the solution structure and dynamics of the RNF4 SIMs region. A series of RNF4 N-terminal (RNF4N) peptides containing pairs of cysteine residues were generated (RNF4N 30/57, 44/70 and 57/84), and subsequently labelled with Cy3B/Alexa647 FRET pair (R_0_~60 Å (16)). Dye labelling did not interfere with SUMO chain binding (Extended data Fig. 2). Using the NMR “kinked” structure (8) of the SIM 2+3 region bound to a SUMO dimer and the Förster distance (R_0_ 60 Å) of the FRET pair, we predicted a FRET efficiency value (E_FRET_) of 0.6 using the accessible volume method (17) (fig. 1B). As a guide, unbound stretched state modelling using a peptide scaffold (derived from PDB: 5M1U) gave an E_FRET_ value of 0.23 (fig. 1B). smFRET histograms of the RNF4N peptides 30/57, 44/70 and 57/84 showed a major population (>90%) with a high-FRET state (E_FRET_ ~0.6-0.7) (fig. 1C), consistent with a kinked structure even in the absence of SUMO chains. The remaining population (~10%) displayed a low-FRET value (E_FRET_ ~0.25-0.3), similar to that predicted for the stretched conformation (fig. 1B). The addition of SUMO chains did not alter significantly the E_FRET_ value (ΔE_FRET_ <0.06) (fig. 1D) or the relative contribution of each population, suggesting the SIMs region already exists in a compact shape primed for engagement with a SUMO polymer. Positioning the FRET dyes outside all four SIMs, RNF4N 37/77, resulted in an E_FRET_ ~ 0.4 (fig. 1 C, Extended data fig. 3). This is indicative of a collapsed conformation as the inter-dye distance in RNF4N 37/77 is such that in an extended conformation it would be beyond FRET detection. Single-molecule FRET trajectories of each RNF4N peptide displayed a single long-lived FRET state with no conformational dynamics, lasting ~10-20 seconds before photobleaching occurred (fig. 1E, Extended data fig. 4-7). SUMO chains did not induce any conformational changes, indicating that the SIMs region is compact, stable, and primed to engage its SUMO chain substrate (fig. 1E, Extended data fig. 4-7).

**Figure 1.**
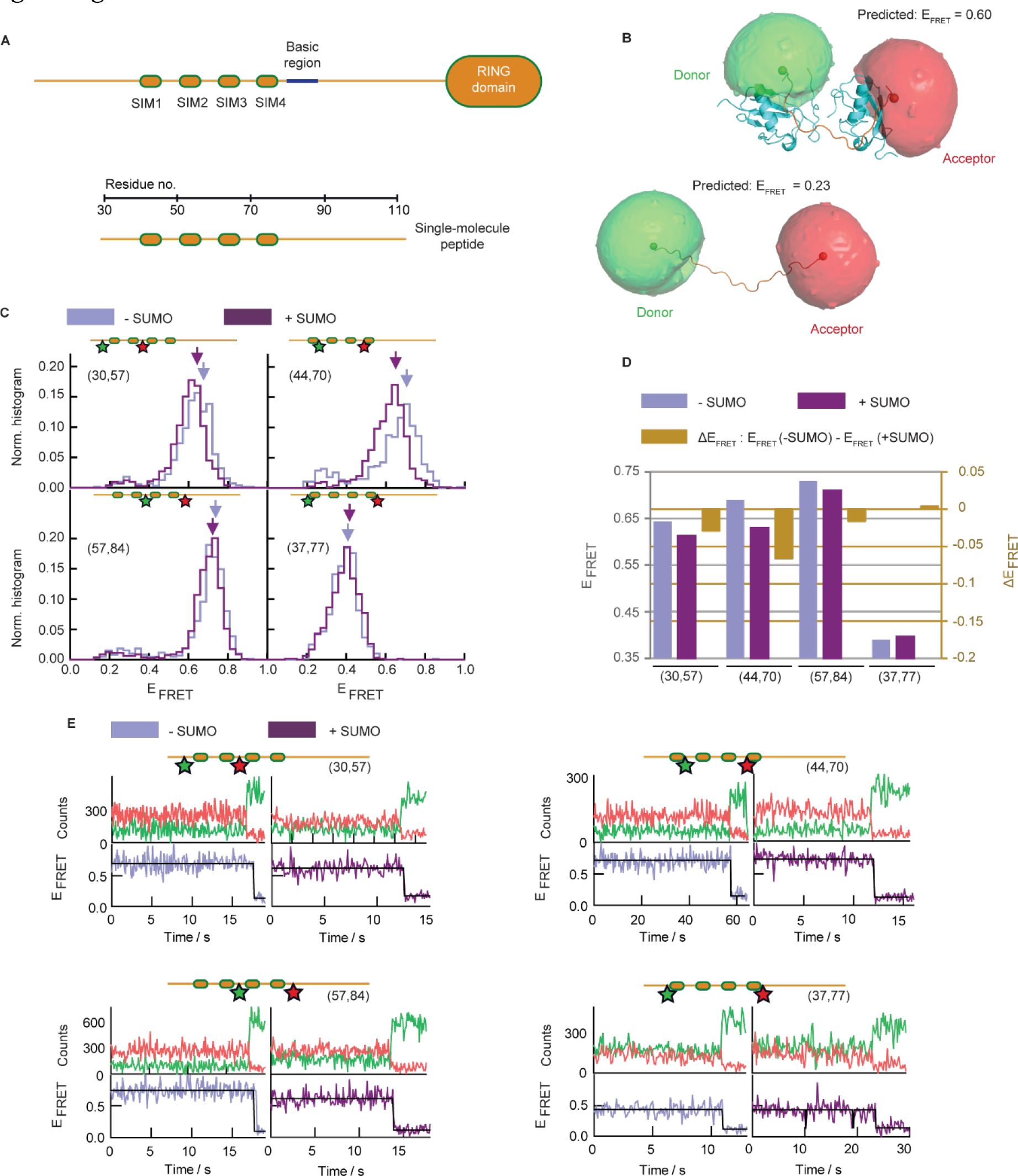
Role of RNF4 N-terminus conformation in substrate binding. **A**, domain layout of RNF4. The unstructured N-terminal region contains four SUMO interaction motifs (SIMs) connected via a linker (~60 residues) to the RING domain. **B top**: NMR structure (PDB: 2MP2) of the SIM2/3 region of RNF4 bound to a SUMO2 dimer (8) modified to include the donor (green) and acceptor (red) fluorophores used for single molecule analysis of RNF4, with a predicted FRET efficiency (E_FRET_) of 0.6. **B bottom**: A modelled peptide (PRB: 5M1U) with similar inter-dye distance but a stretched conformation produced a predicted E_FRET_ value of 0.23. **C**, normalised single-molecule FRET histograms of RNF4N peptides with positions of the donor (green) and acceptor (red) dyes indicated. RNF4N was measured in the absence (light purple) and presence (dark purple) of SUMO chains. Arrows indicate FRET populations used for comparison with/without SUMO (in D). Single-molecule histograms were built from more than 500 molecules. **D**, change in E_FRET_ upon SUMO addition (dark yellow) from C was calculated as: E_FRET_(+SUMO)- E_FRET_(-SUMO). **E**, representative single-molecule trajectories obtained for RNF4N peptides in C. Top panels show donor (green) and acceptor (red) intensity signals. Bottom panels show the E_FRET_ trajectory derived from the donor/acceptor intensity traces in the absence (light purple) and presence (dark purple) of SUMO chains.

The SIMs in RNF4 are short stretches of hydrophobic residues separated by polar residues. To ask if SIM-SIM interactions determined the compact shape, RNF4N^ΔSIMs^ bearing mutations that abolish SUMO binding (Extended data fig. 2) were labelled identically to wild type (WT) RNF4N. Single-molecule FRET histograms of RNF4^ΔSIMs^ showed ~80% of molecules in a high-FRET state (fig. 2A) similar to WT (ΔE_FRET_ <0.03, fig. 2B) but with a slightly higher occupancy of the low-FRET state (E_FRET_ ~0.3) (fig. 2A). smFRET trajectories of RNF4^ΔSIMs^ also showed no evidence of conformational dynamics (Extended data fig. 9-11). The hydrophobic SIMs reside within a highly acidic region of the RNF4 N-terminal domain. Directly adjacent to this within the linker region between SIMs and RING is a cluster of basic residues (fig. 1A). To determine if these residues contributed to the compact shape of RNF4N, three mutants were produced with the basic side chains removed (RNF4N^ΔBasic^) (fig. 2C). During gel filtration RNF4N^ΔBasic^ peptides eluted earlier than their wild type counterparts, suggesting a larger hydrodynamic radius and less compact conformation (Extended data fig. 12A). Labelled RNF4N^ΔBasic^ peptides retained SUMO chain binding (Extended data fig. 12B). smFRET histograms of RNF4N^ΔBasic^ peptides revealed a decrease in E_FRET_ of the major population compared to their WT counterparts (fig. 2C), ranging between 0.08 and 0.19 (fig. 2D, Extended data table 3). The largest ΔE_FRET_ (0.19) was observed for the RNF4N^ΔBasic^ 57/84 peptide reporting the distance between the SIM 2+3 linker and the basic region, contrasting with the RNF4N^ΔSIMs^ 57/84 peptide that displayed a similar FRET profile to WT (fig. 2A, B). The distance increase observed for all RNF4N^ΔBasic^ compared to WT indicates that loss of the basic region induces the RNF4 N-terminal region to “open up” into an extended arrangement.

**Figure 2.**
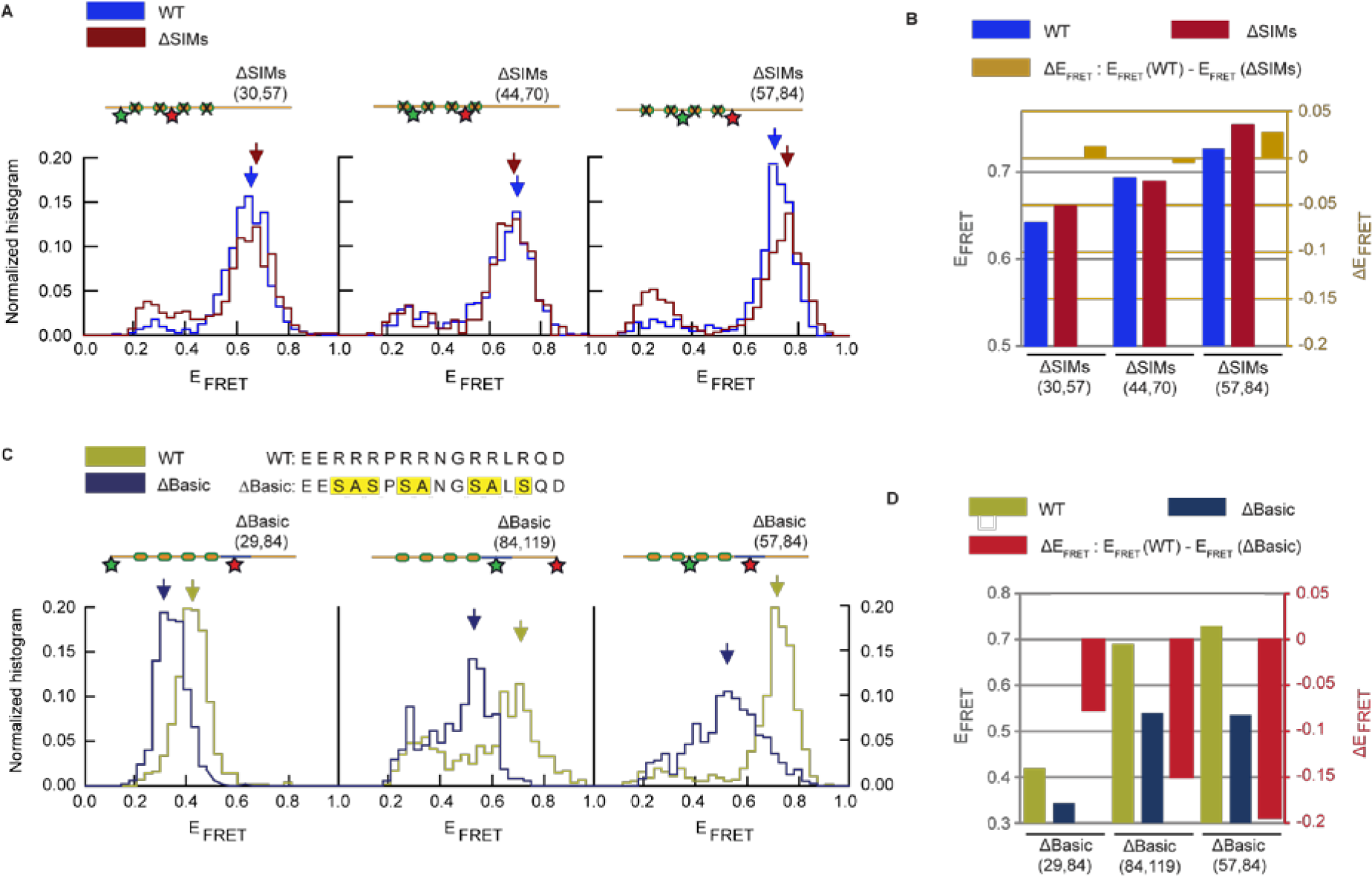
Role of SIMs and basic region in RNF4N conformation. **A**, Single-molecule FRET histograms obtained from RNF4N WT (blue) and ΔSIMs mutant peptides (red) labelled with donor/acceptor FRET dyes (positions indicated inset). **B**, comparison of FRET efficiencies from the main FRET populations (arrows in A) for RNF4N^ΔSIMs^ (red), WT RNF4N (blue) and relative FRET efficiency changes (ΔE_FRET_) between ΔSIM mutants and the corresponding WT (yellow). **C top**, basic region mutations in RNF4N (ΔBasic). **C bottom, s**ingle-molecule FRET histograms from RNF4N WT (yellow) ΔBasic peptides (blue) labelled with donor/acceptor FRET dyes (positions indicated inset). **D**, relative change (red) in FRET efficiency (ΔE_FRET_) between each ΔBasic mutant (blue) and corresponding WT peptide (yellow). Single-molecule FRET histograms shown in A and C are normalised to area unity and built from 400-700 molecules.

To provide insight into the global conformation of RNF4N (fig. 3A) we employed NMR paramagnetic relaxation enhancement (PRE). MTSL spin labels were introduced at positions 29, 44, 58, 70 and 119 of ^15^N-enriched RNF4N (Extended data fig. 17, 18). PREs are observed by comparison of peak intensities from 2D ^1^H-^15^N HSQC spectra measured on MTSL-labelled samples before and after reduction by ascorbate. Relaxation rates for residues located within ~25 Å of the spin label are enhanced and result in peaks with reduced intensity. Comparison of the previously assigned (8) ^1^H-^15^N HSQC spectra of RNF4N before (fig. 3B, lower panel) and after (fig. 3B, upper panel) ascorbate reduction (fig. 3B upper panel) indicated a number of significant intensity changes. PREs are observed for residues 74 and 75 (adjacent to the basic region) in response to a spin label present at either end, indicating that both termini spend a significant proportion of their time within about 25 Å of the central residues. Similarly RNF4N labelled at residues 44, 58 and 70 revealed PREs at one or both termini (fig. 3C). Due to broadening of the peaks from chemical exchange within the SUMO complex, several residues peaks are absent in the spectra. In RNF4N labelled at residue 70 the distance between the spin label and the termini would be over 100 Å in a purely extended conformation, therefore PRE determinations indicate that both termini fold back towards the middle of the sequence.

**Figure 3.**
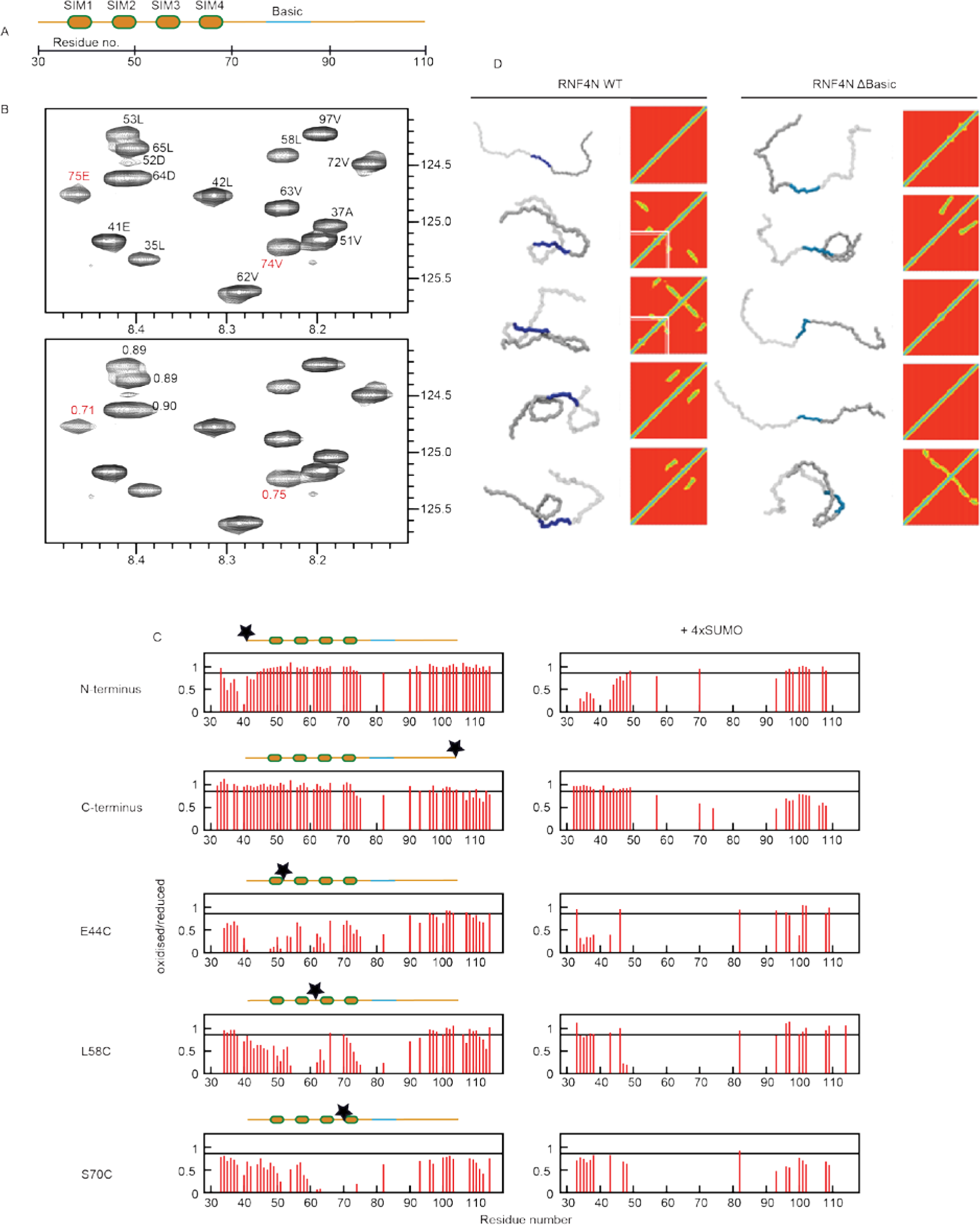
NMR and molecular dynamic simulations reveal compact shape of RNF4. **A**, RNF4N peptide layout used in B. **B top**, reference ^1^H-^15^N HSQC spectrum of reduced RNF4 using peak assignment established previously (8). **B bottom,**same region prior to MTSL reduction. Peaks with significant ratio changes adjacent to the basic region, 74E and 75V (red), indicated. **C**, histograms of HSQC peak intensity ratios plotted for RNF4N labelled at the termini, E44C, L58C, and S70C (left) and in complex with SUMO chains (right). Peaks are absent due to severe peak broadening in the SUMO complex. Significant intensity reductions are below 0.85 (black line). **D**, molecular dynamic simulations of RNF4N WT (left) and ΔBasic mutant (right). Top representative conformations obtained by conformational clustering analysis of five independent simulations. Polypeptide backbone is transparent grey for residues 32-76, blue for basic region residues 77-86, and dark grey for residues 87 to 133. Contact plots show residue-residue (X-Y axis) contacts for each cluster. Electrostatic interactions involving the basic region are highlighted in simulations 2 and 3 by white lines. Properties calculated from simulations indicated in Extended data table 4.

To capture potential conformations of RNF4 in its disordered state, we used unbiased coarse-grained (18) molecular dynamics simulations, in which an initial fully-extended model peptide of RNF4 residues 32-133 was simulated in solution. The top clusters (19) of conformations from each of five independent trajectories and their corresponding residue-by-residue contact plots indicate that the RNF4 N-terminal peptide adopts both flexible and compact conformations (fig. 3D). The contact plot for these conformers highlights the basic region between R77 and R88 suggesting that electrostatic interactions involving these residues play a prominent role in driving compaction (fig. 3D). Consistent with smFRET and NMR data, the RNF4N^ΔBasic^ mutant tends to occupy more extended conformations (fig. 3D), highlighted by measuring the distance from residue 80 to 133 (Extended data Table 4). Absent from the contact plots of RNF4N^ΔBasic^ is an interaction between the mutated basic region and the N-terminal half of the protein, indicated by white lines in the plots (fig. 3D, Extended figure 19).

To address the role of the basic region in the function of RNF4 as a SUMO targeted ubiquitin E3 ligase we compared the ability of full length versions of RNF4 and RNF4^ΔBasic^ to incorporate fluorescently labelled ubiquitin in response to increasing 4xSUMO substrate concentrations. In the presence of 4xSUMO substrate, RNF4 catalysed the incorporation of free ubiquitin into higher molecular weight material and this was severely compromised with RNF4^ΔBasic^ (fig. 4A, B). To compare the rates of ubiquitination, a real-time fluorescence polarization (FP) assay was performed to monitor the incorporation of fluorescent ubiquitin into higher molecular weight products (fig. 4C). The initial rates showed that both RNF4 and RNF4^ΔBasic^ display a lag phase before incorporation of ubiquitin accelerates. For RNF4^ΔBasic^ this lag phase is extended and the rate of incorporation remains slower (fig. 4C). Comparing the modification rate of fluorescently labelled 4xSUMO, RNF4^ΔBasic^ had a 3-fold reduced initial rate of ubiquitination (fig. 4D, E). Although the basic region is more than forty residues from the RING domain, it was important to establish its influence on intrinsic RING activity. This was evaluated in a substrate independent lysine discharge assay where free lysine acts as the nucleophile and attacks the thioester bond between ubiquitin and E2, releasing free ubiquitin. A functional E3 ligase activates the thioester bond rendering it susceptible to nucleophilic attack by lysine. Both RNF4 and RNF4^ΔBasic^ released ubiquitin at similar rates (fig. 4F, G), indicating that the mutations did not affect the intrinsic catalytic activity of the RING. Differences in ubiquitination activity are not due to differences in substrate binding as RNF4 and RNF4^ΔBasic^ bind SUMO chains with similar affinities (fig. 4H).

**Figure 4.**
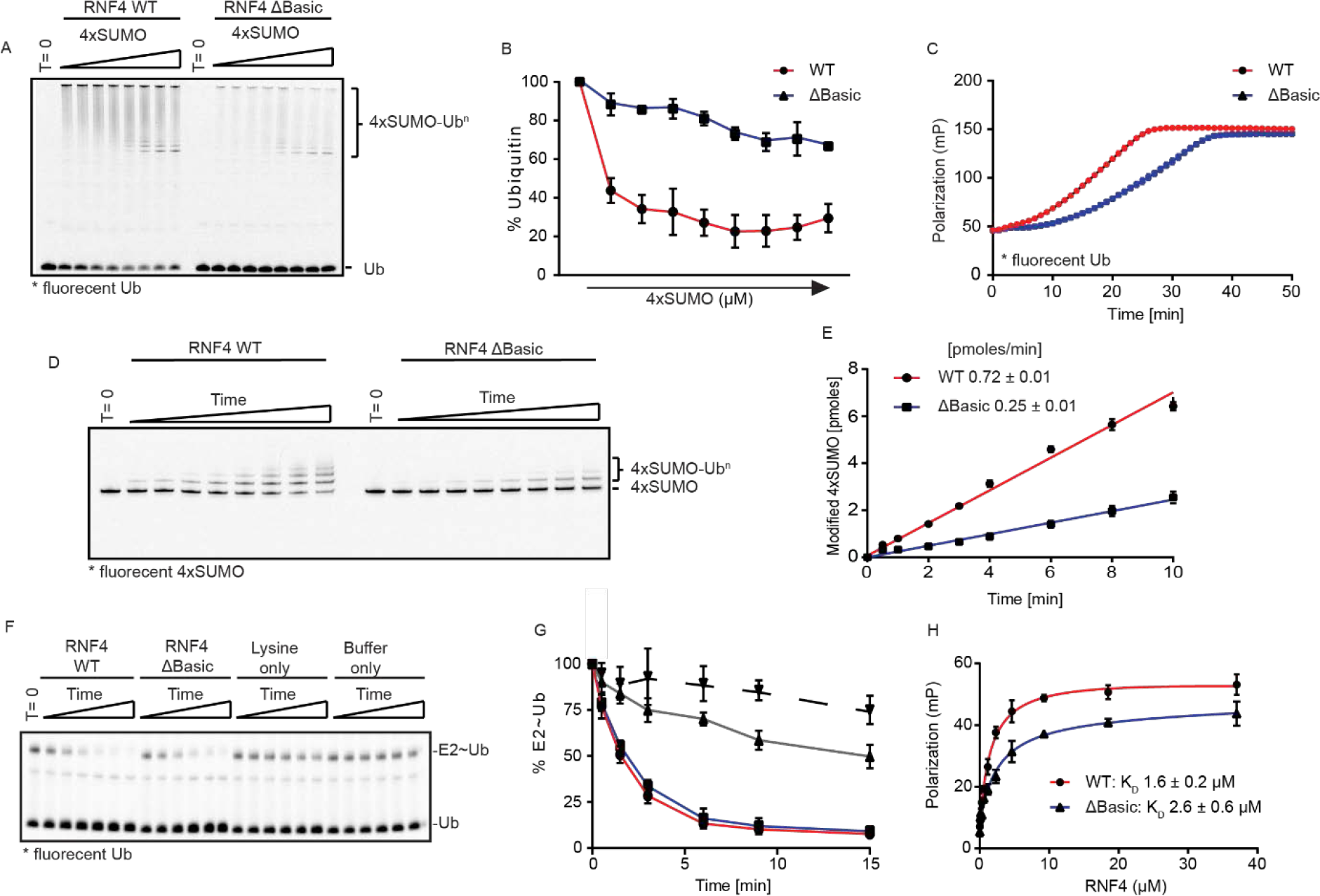
Basic region mutant displays reduced ubiquitination activity. **A**, ubiquitination activity of RNF4 wild type (WT) and ΔBasic mutant in response to increasing substrate (4xSUMO) concentrations. Assays (15 min) contained E1 activating enzyme, UbcH5a and fluorescently labelled ubiquitin. T = 0 is without ATP. Reaction products were fractionated by SDS-PAGE, followed by in-gel fluorescence. **B**, quantitation of fluorescent free ubiquitin in A (WT red, ΔBasic blue). Ubiquitin levels at T = 0 represented 100%. **C**, fluorescence polarization real-time ubiquitination assay comparing activity of WT (red) against ΔBasic (blue). Assays contained 4xSUMO, E1, UbcH5a and fluorescent ubiquitin. Reactions initiated with ATP addition and FP measured over 50 minutes. **D**, substrate ubiquitination activity assay containing fluorescently labelled 4xSUMO, ubiquitin, E1, UbcH5a and either RNF4 WT or RNF4 ΔBasic. Reaction products analysed as in A. **E**, quantitation of D for RNF4 WT (red) and RNF4 ΔBasic (blue). **F**, lysine discharge assays using fluorescently labelled ubiquitin. The discharge of ubiquitin from the E2 (E2~Ub) was analysed by SDS-PAGE over time in the presence of RNF4 WT, RNF4 ΔBasic, lysine only, or buffer only. **G**, measurement of fluorescence from the E2~Ub as seen in F. T = 0 represents 100% E2~Ub (WT, red; ΔBasic, blue; lysine only, grey; buffer only, dashed). **H**, binding affinity of RNF4 for 4xSUMO via FP. A fixed 4xSUMO concentration was titrated with RNF4 (WT, red; ΔBasic, blue) to saturating concentrations. For each graph (B, C, E, G, H) data represent mean ± standard deviation (n = 3).

The substrate adaptor region of RNF4 contains multiple SUMO Interaction Motifs (SIMs) embedded in what was thought to be an intrinsically disordered region. However our studies have revealed a global three-dimensional architecture within this “disordered” region, with highly flexible but compact structures facilitating productive binding of SUMO substrate. Located between the SIMs and the RING is a basic linker that mediates delivery of substrate to the catalytically primed RING-E2~Ubiquitin. As many ubiquitin E3 ligases or their targets contain regions shown or predicted to be disordered (20–22), this work provides a framework for understanding how the RING and substrate adapter regions of an E3 ligase co-operate to ensure efficient ubiquitin transfer. More generally it provides new insight into how dynamic and intrinsically disordered proteins can adopt ensembles containing preferred structures that are essential for function, and supports the idea of electrostatics driving intermolecular interactions between disordered proteins (23).

## Materials and Methods

### Cloning and production of recombinant proteins

All constructs are described in Extended data tables 5 and 6. Full length RNF4 variants (genes synthesized by GenScript) were expressed from the pLOU3 vector in Escherichia coli BL21 (DE3) cells, while RNFN 27-118 variants (gene fragments produced as gBlocks by IDT and recombined into pHis-TEV-30a using New England BioLabs HiFi assembly) were expressed from pHis-TEV-30a vector in E. coli Rosetta (DE3) cells as previously described (1,2). 15N-enriched RNF4N 30-118 and 32-133 (gBlocks recombined into pHis-TEV-30a using New England BioLabs HiFi assembly) were expressed from pHis-TEV-30a vector in E. coli Rosetta (DE3) cells as previously described (3). For 15N-enriched peptides cells were grown at 37°C in M9 minimal medium supplemented with 15NH4Cl (Sigma). Samples were purified by Ni-NTA (Qiagen) chromatography following cleavage with TEV protease. RNF4N peptides were further purified to homogeneity by gel filtration (following biotinylation for single-molecule peptides).

Linear SUMO2 (genes synthesized by GenScript) genes bearing C47A mutations (4xSUMO) was subcloned into pHis-TEV-30a vector, along with a second variant containing a single cysteine at the C-terminus. Fusion proteins were purified as described for RNF4N peptides.

### Labelling proteins/peptides with functional groups

RNF4N peptides for single-molecule measurements were biotinylated within a C-terminal AviTag using the E. coli biotin ligase BirA (4). Reactions contained 200 μM AviTag-fused RNF4N peptide, 5 mM MgCl2, 200 mM KCl, 2.5 mM ATP, 800 μM D-biotin in 50 mM Tris, 150 mM NaCl buffer at pH 7.5 (total volume 500 μl). Reactions were incubated at room temperature for 4 hours, followed by overnight incubation at 4°C. Biotinylated RNF4 was then purified by gel filtration and correct mass confirmed by LCMS (see example Extended data figure 1). Next, RNF4N peptides were labelled stochastically with FRET dyes at a 2:2:1 molar ratio (Cy3B: Alexa647: RNF4N), in 50 mM Tris, 150 mM NaCl at pH 7. FRET dyes used contained maleimide function groups for cysteine labelling, with dyes dissolved in DMSO (Cy3B, GE Healthcare; Alexa647, Invitrogen). Reactions were incubated at room temperature for 2 hours before filtration to remove unreacted dyes using Centripure P2 filtration columns, into 50 mM Tris, 150 mM NaCl at pH 7.5. Correct mass of labelled peptides was confirmed by LCMS (see example Extended data figure 1). 4xSUMO fusion protein containing a single cysteine was labelled with Alexa488 (Invitrogen) as described for RNF4N.

15N-enriched RNF4N peptides containing a single cysteine for MTSL labelling were stored under reducing conditions in 50 mM Tris, 150 mM NaCl, 0.5 mM TCEP at pH 7.5. Immediately prior to labelling reaction, peptides were buffer exchanged using a Centripure P2 filtration column into 50 mM Tris, 150 mM NaCl at pH 7.4. Reactions were performed at a 5:1 molar ratio of MTSL (dissolved in acetonitrile) over RNF4N for 1 hour at room temperature. Unreacted MTSL was removed using a Centripure P2 filtration column into 50 mM Tris, 150 mM NaCl at pH 7.4, with correct mass of labelled peptide confirmed by LCMS.

### Native gel binding assay

Due to the stochastic labelling of RNF4N peptides with FRET dyes, final concentration were only approximations. To assess binding to 4xSUMO (or 6xSUMO) native gel binding assays were performed (5). RNF4N was mixed with 4xSUMO, final concentrations ~2 μM and 10 μM respectively, and incubated at room temperature for 10 minutes in a total volume of 20 μl. Protein binding reactions were performed in 50 mM Tris, 150 mM NaCl, 0.5 mM TCEP at pH 7.5. Next, native loading buffer was added to samples resulting in a 10% (v/v) glycerol concentration, before loading onto 6% DNA retardation gels (Invitrogen). Samples were separated 60 V, 4°C in 0.5x tris-borate-EDTA buffer before analysis by in-gel fluorescence using a Bio-Rad ChemiDoc Imaging System.

### Molecular modelling of dye positions

For the three-dimensional FRET models produced, the NMR structure PDB: 2MP2 (6) was used as template for the “kinked” model, while the NMR structure PDB: 5M1U (7) was used as template for the extended model. The modelling of dyes was performed using FRET positioning software developed by the Seidel lab (8). The software produces a sphere that approximates each dye defined by three radii, attached to the peptide by a flexible linker defined by its length (Llinker) and width (Wlinker). For Cy3B: Llinker 20.5 Å, Wlinker 4.5 Å, Rdye1 6.8 Å, Rdye2 3.0 Å, Rdye3 1.5 Å. For Alexa647: Llinker 21 Å, Wlinker 4.5 Å, Rdye1 11 Å, Rdye2 4.7 Å, Rdye3 1.5 Å. The parameters are used by the software to calculate the dye accessible volume (AV) that simulates all accessible positions through manipulation of the linker from the point of attachment to the peptide. The attachment positions used for the “kinked” structure on PDB: 2MP2 are four residues shorter than those used in RNF4N 44/70 (23 and 27 respectively), due to limited access to residue sides chains with the structure. The attachment positions used for the extended structure PDB: 5M1U have the same residue separation as RNF4N 44/70. The modelled dyes were then used to produce predicted FRET efficiency values. This prediction uses the R0 of 60 Å previously reported for this FRET pair (9).

### Single-molecule total-internal reflection

Single-molecule FRET (smFRET) experiments were performed as previously reported (10). smFRET intensity traces were acquired from immobilized peptides using a prism-type total-internal reflection setup that includes an inverted microscope (Olympus IX71) coupled to a 532-nm laser (Crystalaser) and a back illuminated Ixon EMCCD camera (Andor). Donor and acceptor fluorescence intensities were separated using dichroic mirrors (DCRLP645, Chroma Technology) and imaged onto the left (donor) and right (acceptor) half-chip of the EMCCD. This setup allowed us to monitor the Cy3B and the Alexa647 signals simultaneously. Images were processed using IDL and data analysis was performed using laboratory-written routines in Matlab as previously described (11). smFRET trajectories were acquired at 100 ms integration time unless stated otherwise. FRET was calculated from the raw donor and acceptor intensity traces using EFRET = IA/(ID + αIA) where a =0.88 accounts for 12% leakage into the acceptor detection channel (11). When constructing FRET population histograms, the first ten frames of each movie were averaged to produce an ‘average’ FRET value for that movie. Data were acquired in imaging buffer (50 mM Tris–HCl (pH 7.5), 150 mM NaCl, 6% (w/w) glucose, 0.1 mg/ml glucose oxidase (Sigma) and 0.02 mg/ml glucose catalase (Sigma), 1 mM Trolox.

### Paramagnetic relaxation enhancement (PRE) NMR spectroscopy

A final concentration of ~150 μM 15N-enriched RNF4N fragments were used for NMR measurements, and SUMO chains were added for a monomer ratio of 1:2 for RNF4N:SUMO in the complex studies. Standard 1H-15N HSQC spectra were recorded for these MTSL labelled samples, which were then reduced by adding 2 mM sodium ascorbate and incubated at 4 °C overnight, and 1H-15N HSQC were recorded using the same settings on reduced samples. All experiments were run on a Bruker Avance HD III 800MHz NMR spectrometer at 25 °C. The intensity ratio were then converted into distance restraints (12).

### Molecular dynamics simulations

Initial fully extended conformations of RNF4N wildtype and basic mutant were generated using Modeller v9 (13) and processed with the PDB2PQR server (14). These were then mapped to course-grained (CG) using SIRAH Tools (15) The CG models were solvated using pre-equilibrated WT4 molecules in octahedral boxes of 2.0 nm size from the solute. An ionic strength of 0.15 M was set by randomly replacing WT4 molecules by Na+ and Cl- CG ions. The system was prepared following standard SIRAH protocols: 5000 steps of steepest descent energy minimisation with positional restraints of 1000 kJ mol-1 nm-2 on backbone particles, followed by 5000 steps with no backbone restraints, followed by 5 ns NVT simulation at 300 K with 1000 kJ mol-1 nm-2 restraints on the protein to equilibrate the solvent, and a 25 ns simulation with reduced positional restraints of 100 kJ mol-1 nm-2 on backbone beads to improve side chain solvation. Production simulations used the NPT ensemble at 300 K and 1 bar. Simulations were performed using GROMACS v5 (http://www.gromacs.org) (16). PME electrostatics were used with the Verlet scheme with a cut off of 1.2 nm. Solvent and solute were coupled separately to V-rescale thermostats (17) with coupling times of 2 ps. Pressure was controlled by the Parrinello-Rahman barostat (18) with a coupling time of 8 ps. Production trajectories were performed for. Trajectory analysis and visualisation used VMD (19) and gromacs tools. Residue-residue contact plots were generated using the mdmat tool for Cα atoms.

### Ubiquitination assays

Ubiquitination assays were performed as described previously (20). To measure total ubiquitination assays contained fluorescently labelled ubiquitin (Fluorescein, Invitrogen). Reactions were carried out by mixing the following components: 0.2 μM E1, 1 μM UbcH5a, 0.4 μM RNF4, 9 μM ubiquitin, 1 μM ubiquitin-Fluorescein, 3 mM ATP, 5 mM MgCl2, 50 mM Tris, 150 mM NaCl, 0.5 mM TCEP. 4xSUMO substrate titration range: 0.125 – 8 μM. Reactions were quenched with SDS-PAGE loading buffer. Samples were subject to SDS-PAGE electrophoresis before analysis for in-gel fluorescence using a Bio-Rad ChemiDoc Imaging System. T=0 samples were taken before the addition of ATP.

Direct 4xSUMO substrate ubiquitination assays were performed as described above with the following modifications. Ubiquitin was unlabelled and added at 20 μM. The 4xSUMO substrate was added at 1 μM, with 4xSUMO-Alexa488 at 0.25 μM.

Real-time ubiqutination assay were performed as described previously (21) on a PHERAstar (BMG labtech) via an FP optics module (FP 485 520). Assays were performed in a Greiner microplate (384 well, black, non-binding) and measured over the course of 50 minutes in a total volume of 20 μl. Reactions were carried out by mixing the following components: 0.1 μM E1, 1 μM UbcH5a, 0.4 μM RNF4, 9.6 μM ubiquitin, 0.4 μM ubiquitin-Fluorescein, 3 μM 4xSUMO, 3 mM ATP, 5 mM MgCl2, 50 mM Tris, 150 mM NaCl, 0.5 mM TCEP.

### Lysine discharge assay

Lysine discharge assays were performed as previously described (20). E2 (Ubch5a) was first loaded with ubiquitin in the absence of a substrate or E3. The E2~Ub thioester was prepared by incubating 100 μM E2 with 114 μM ubiquitin and 6 μM ubiquitin-Fluorescein, along with 0.2 μM E1 in 50 mM Tris, 150 mM NaCl, 3 mM ATP, 5 mM MgCl2, 0.1% NP40 at pH 7.5. The reaction was incubated at 37°C for 12 minutes. To stop the loading of E2 with ubiquitin by E1, ATP was depleted using Apyrase (4.5 U/ml; New England BioLabs) at room temperature for 10 minutes. The E2~Ub thioester was then mixed at a 1:1 ratio with RNF4 and L-lysine in 50 mM Tris, 150 mM NaCl at pH 7.5 (final concentrations at 0.3 μM RNF4 and 25 mM L-lysine). The Reactions were incubated at room temperature before quenching using non-reducing SDS-PAGE loading buffer. Samples were subject to SDS-PAGE electrophoresis before analysis for in-gel fluorescence using a Bio-Rad ChemiDoc Imaging System. T=0 was taken before the E2~Ub thioester was mixed with RNF4 and L-lysine

### Affinity assay by fluorescence polarization

The binding affinity between RNF4 and 4xSUMO was performed via fluorescence polarization with settings as described for real-time ubiquitination assay. Unlabelled RNF4 was titrated against 1 μM 4xSUMO-Alexa488 in 50 mM Tris, 150 mM NaCl, 0.5 mM TCEP at pH 7.5. Reactions were incubated at room temperature for 10 minutes before measuring FP.

## Acknowledgements

This work was supported by the following grants: Wellcome Trust Investigator Award (098391/Z/12/Z) to R.T.H., Wellcome Trust Studentship (109113/Z/15/Z) to P.M., Wellcome Trust Collaborative Award (215539) and multiuser equipment grant (104833) to S.J.M. Additionally J.C.P. thanks the Scottish Universities Physics Alliance (SUPA) and the University of St. Andrews for financial support.

## Author contributions

P.M. cloned expressed and purified proteins, conducted smFRET, smFRET modelling, biochemical experiments and performed data analysis. J.C.P. conducted smFRET experiments and performed data analysis. S.J.M. designed NMR experiments and contributed to data analysis. Y.X. conducted NMR experiments and performed data analysis. S.L.R. conducted molecular simulations and performed data analysis. R.T.H. conceived the project and contributed to data analysis. All authors contributed to manuscript preparation.

## Data availability

Raw data are available upon request to the authors.

## References

1. Deshaies, R. J., and Joazeiro, C. A. (2009) RING domain E3 ubiquitin ligases. Annual review of biochemistry 78, 399–434

2. Plechanovova, A., Jaffray, E. G., McMahon, S. A., Johnson, K. A., Navratilova, I., Naismith, J. H., and Hay, R. T. (2011) Mechanism of ubiquitylation by dimeric RING ligase RNF4. Nature structural & molecular biology 18, 1052–1059

3. Plechanovova, A., Jaffray, E. G., Tatham, M. H., Naismith, J. H., and Hay, R. T. (2012) Structure of a RING E3 ligase and ubiquitin-loaded E2 primed for catalysis. Nature 489, 115–120

4. Tatham, M. H., Geoffroy, M. C., Shen, L., Plechanovova, A., Hattersley, N., Jaffray, E. G., Palvimo, J. J., and Hay, R. T. (2008) RNF4 is a poly-SUMO-specific E3 ubiquitin ligase required for arsenic-induced PML degradation. Nature cell biology 10, 538–546

5. Lallemand-Breitenbach, V., Jeanne, M., Benhenda, S., Nasr, R., Lei, M., Peres, L., Zhou, J., Zhu, J., Raught, B., and de The, H. (2008) Arsenic degrades PML or PML-RARalpha through a SUMO-triggered RNF4/ubiquitin-mediated pathway. Nature cell biology 10, 547–555

6. Galanty, Y., Belotserkovskaya, R., Coates, J., and Jackson, S. P. (2012) RNF4, a SUMO-targeted ubiquitin E3 ligase, promotes DNA double-strand break repair. Genes & development 26, 1179–1195

7. Yin, Y., Seifert, A., Chua, J. S., Maure, J. F., Golebiowski, F., and Hay, R. T. (2012) SUMO-targeted ubiquitin E3 ligase RNF4 is required for the response of human cells to DNA damage. Genes & development 26, 1196–1208

8. Xu, Y., Plechanovova, A., Simpson, P., Marchant, J., Leidecker, O., Kraatz, S., Hay, R. T., and Matthews, S. J. (2014) Structural insight into SUMO chain recognition and manipulation by the ubiquitin ligase RNF4. Nature communications 5, 4217

9. Song, J., Durrin, L. K., Wilkinson, T. A., Krontiris, T. G., and Chen, Y. (2004) Identification of a SUMO-binding motif that recognizes SUMO-modified proteins. Proceedings of the National Academy of Sciences of the United States of America 101, 14373–14378

10. Hecker, C. M., Rabiller, M., Haglund, K., Bayer, P., and Dikic, I. (2006) Specification of SUMO1- and SUMO2-interacting motifs. The Journal of biological chemistry 281, 16117–16127

11. Dou, H., Buetow, L., Sibbet, G. J., Cameron, K., and Huang, D. T. (2012) BIRC7-E2 ubiquitin conjugate structure reveals the mechanism of ubiquitin transfer by a RING dimer. Nature structural & molecular biology 19, 876–883

12. Pruneda, J. N., Littlefield, P. J., Soss, S. E., Nordquist, K. A., Chazin, W. J., Brzovic, P. S., and Klevit, R. E. (2012) Structure of an E3:E2~Ub complex reveals an allosteric mechanism shared among RING/U-box ligases. Molecular cell 47, 933–942

13. Scott, D. C., Sviderskiy, V. O., Monda, J. K., Lydeard, J. R., Cho, S. E., Harper, J. W., and Schulman, B. A. (2014) Structure of a RING E3 trapped in action reveals ligation mechanism for the ubiquitin-like protein NEDD8. Cell 157, 1671–1684

14. Branigan, E., Plechanovova, A., Jaffray, E. G., Naismith, J. H., and Hay, R. T. (2015) Structural basis for the RING-catalyzed synthesis of K63-linked ubiquitin chains. Nature structural & molecular biology 22, 597–602

15. Streich, F. C., Jr., and Lima, C. D. (2016) Capturing a substrate in an activated RING E3/E2-SUMO complex. Nature 536, 304–308

16. Chakraborty, A., Wang, D., Ebright, Y. W., Korlann, Y., Kortkhonjia, E., Kim, T., Chowdhury, S., Wigneshweraraj, S., Irschik, H., Jansen, R., Nixon, B. T., Knight, J., Weiss, S., and Ebright, R. H. (2012) Opening and closing of the bacterial RNA polymerase clamp. Science 337, 591–595

17. Kalinin, S., Peulen, T., Sindbert, S., Rothwell, P. J., Berger, S., Restle, T., Goody, R. S., Gohlke, H., and Seidel, C. A. (2012) A toolkit and benchmark study for FRET-restrained high-precision structural modeling. Nature methods 9, 1218–1225

18. Darre, L., Machado, M. R., Brandner, A. F., Gonzalez, H. C., Ferreira, S., and Pantano, S. (2015) SIRAH: a structurally unbiased coarse-grained force field for proteins with aqueous solvation and long-range electrostatics. Journal of chemical theory and computation 11, 723–739

19. Daura, X., Gademann, K., Jaun, B., Seebach, D., van Gunsteren, W. F., and Mark, A. E. (1999) Peptide Folding: When Simulation Meets Experiment. Angewandte Chemie International Edition 38, 236–240

20. Bhowmick, P., Pancsa, R., Guharoy, M., and Tompa, P. (2013) Functional diversity and structural disorder in the human ubiquitination pathway. PloS one 8, e65443

21. Mittag, T., Marsh, J., Grishaev, A., Orlicky, S., Lin, H., Sicheri, F., Tyers, M., and Forman-Kay, J. D. (2010) Structure/function implications in a dynamic complex of the intrinsically disordered Sic1 with the Cdc4 subunit of an SCF ubiquitin ligase. Structure 18, 494–506

22. Pierce, W. K., Grace, C. R., Lee, J., Nourse, A., Marzahn, M. R., Watson, E. R., High, A. A., Peng, J., Schulman, B. A., and Mittag, T. (2016) Multiple Weak Linear Motifs Enhance Recruitment and Processivity in SPOP-Mediated Substrate Ubiquitination. Journal of molecular biology 428, 1256–1271

23. Borgia, A., Borgia, M. B., Bugge, K., Kissling, V. M., Heidarsson, P. O., Fernandes, C. B., Sottini, A., Soranno, A., Buholzer, K. J., Nettels, D., Kragelund, B. B., Best, R. B., and Schuler, B. (2018) Extreme disorder in an ultrahigh-affinity protein complex. Nature 555, 61–66

## References

1. Bruderer, R., Tatham, M. H., Plechanovova, A., Matic, I., Garg, A. K., and Hay, R. T. (2011) Purification and identification of endogenous polySUMO conjugates. EMBO reports 12, 142–148

2. Tatham, M. H., Geoffroy, M. C., Shen, L., Plechanovova, A., Hattersley, N., Jaffray, E. G., Palvimo, J. J., and Hay, R. T. (2008) RNF4 is a poly-SUMO-specific E3 ubiquitin ligase required for arsenic-induced PML degradation. Nature cell biology 10, 538–546

3. Battiste, J. L., and Wagner, G. (2000) Utilization of site-directed spin labeling and high-resolution heteronuclear nuclear magnetic resonance for global fold determination of large proteins with limited nuclear overhauser effect data. Biochemistry 39, 5355–5365

4. Fairhead, M., and Howarth, M. (2015) Site-specific biotinylation of purified proteins using BirA. Methods in molecular biology 1266, 171–184

5. Tatham, M. H., Kim, S., Jaffray, E., Song, J., Chen, Y., and Hay, R. T. (2005) Unique binding interactions among Ubc9, SUMO and RanBP2 reveal a mechanism for SUMO paralog selection. Nature structural & molecular biology 12, 67–74

6. Xu, Y., Plechanovova, A., Simpson, P., Marchant, J., Leidecker, O., Kraatz, S., Hay, R. T., and Matthews, S. J. (2014) Structural insight into SUMO chain recognition and manipulation by the ubiquitin ligase RNF4. Nature communications 5, 4217

7. Schubeis, T., Spehr, J., Viereck, J., Kopping, L., Nagaraj, M., Ahmed, M., and Ritter, C. (2018) Structural and functional characterization of the Curli adaptor protein CsgF. FEBS letters 592, 1020–1029

8. Kalinin, S., Peulen, T., Sindbert, S., Rothwell, P. J., Berger, S., Restle, T., Goody, R. S., Gohlke, H., and Seidel, C. A. (2012) A toolkit and benchmark study for FRET-restrained high-precision structural modeling. Nature methods 9, 1218–1225

9. Chakraborty, A., Wang, D., Ebright, Y. W., Korlann, Y., Kortkhonjia, E., Kim, T., Chowdhury, S., Wigneshweraraj, S., Irschik, H., Jansen, R., Nixon, B. T., Knight, J., Weiss, S., and Ebright, R. H. (2012) Opening and closing of the bacterial RNA polymerase clamp. Science 337, 591–595

10. Blouin, S., Craggs, T. D., Lafontaine, D. A., and Penedo, J. C. (2009) Functional studies of DNA-protein interactions using FRET techniques. Methods in molecular biology 543, 475–502

11. McCluskey, K., Shaw, E., Lafontaine, D. A., and Penedo, J. C. (2014) Single-molecule fluorescence of nucleic acids. Methods in molecular biology 1076, 759–791

12. Sjodt, M., and Clubb, R. T. (2017) Nitroxide Labeling of Proteins and the Determination of Paramagnetic Relaxation Derived Distance Restraints for NMR Studies. Bio-protocol 7

13. Sali, A., and Blundell, T. L. (1993) Comparative protein modelling by satisfaction of spatial restraints. Journal of molecular biology 234, 779–815

14. Dolinsky, T. J., Czodrowski, P., Li, H., Nielsen, J. E., Jensen, J. H., Klebe, G., and Baker, N. A. (2007) PDB2PQR: expanding and upgrading automated preparation of biomolecular structures for molecular simulations. Nucleic Acids Research 35, W522–W525

15. Machado, M. R., and Pantano, S. (2016) SIRAH tools: mapping, backmapping and visualization of coarse-grained models. Bioinformatics 32, 1568–1570

16. Abraham, M. J., Murtola, T., Schulz, R., Páll, S., Smith, J. C., Hess, B., and Lindahl, E. (2015) GROMACS: High performance molecular simulations through multi-level parallelism from laptops to supercomputers. SoftwareX 1–2, 19–25

17. Bussi, G., Donadio, D., and Parrinello, M. (2007) Canonical sampling through velocity rescaling. The Journal of Chemical Physics 126, 014101

18. Parrinello, M., and Rahman, A. (1981) Polymorphic transitions in single crystals: A new molecular dynamics method. Journal of Applied Physics 52, 7182–7190

19. Humphrey, W., Dalke, A., and Schulten, K. (1996) VMD: Visual molecular dynamics. Journal of Molecular Graphics 14, 33–38

20. Branigan, E., Plechanovova, A., Jaffray, E. G., Naismith, J. H., and Hay, R. T. (2015) Structural basis for the RING-catalyzed synthesis of K63-linked ubiquitin chains. Nature structural & molecular biology 22, 597–602

21. Branigan, E., Plechanovova, A., and Hay, R. T. (2019) Methods to analyze STUbL activity. Methods in enzymology 618, 257–280

