## Supplementary figures for "Functional 3D architecture in an intrinsically disordered E3 ligase domain facilitates ubiquitin transfer"

### Extended data

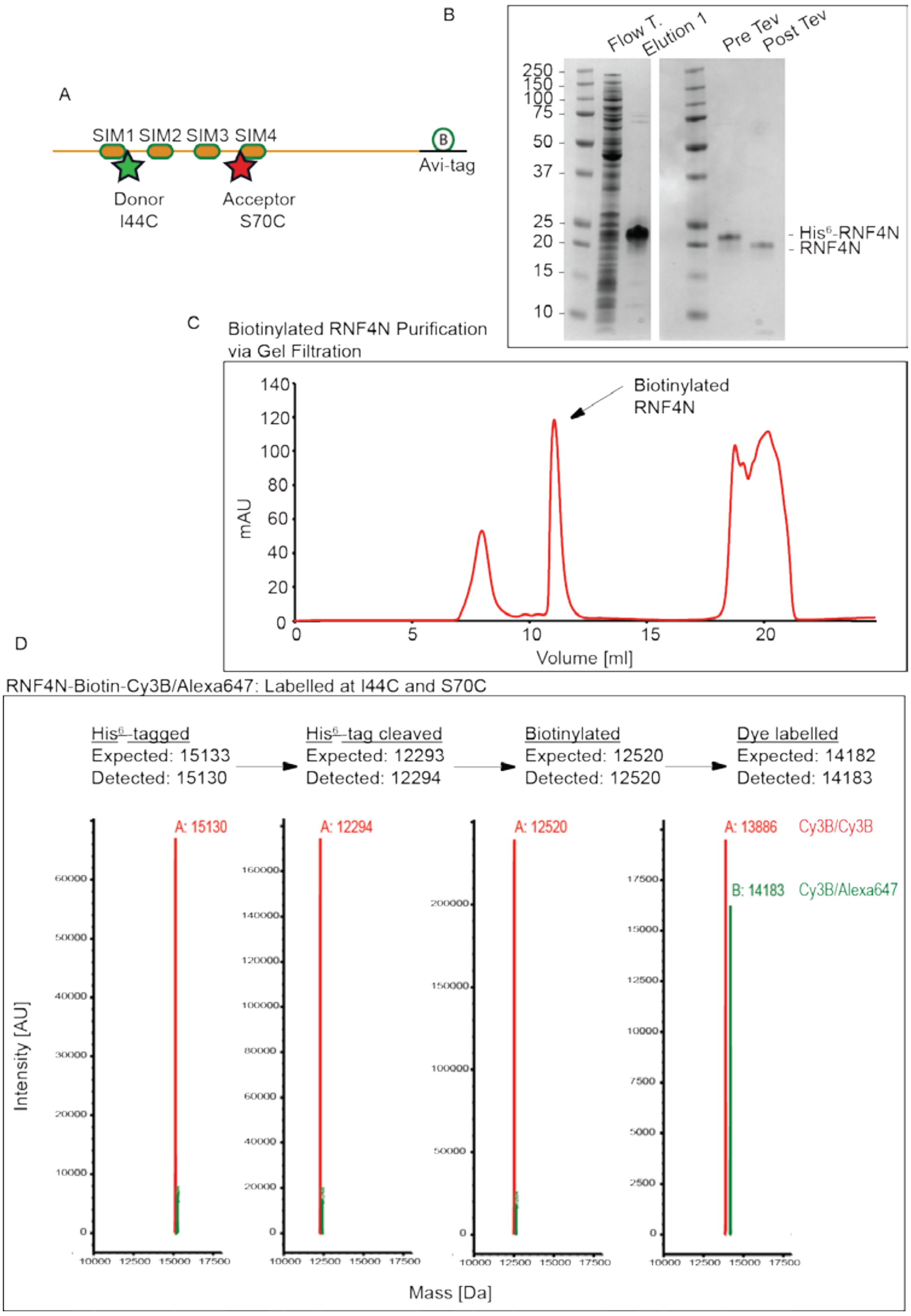

**Extended data figure 1. Production of RNF4N 44/70 peptide.**

A, model of the N-terminal region of RNF4 bearing a C-terminal AviTag (RNF4N). The AviTag allows for site directed biotinylation of and internal lysine residue. The four SIMs are shown, along with the attachment position of the FRET fluorophore pair (residues 40, 70). B, RNF4N was initially expressed with an N-terminal His$^6$-tag to allow for purification. The His$^6$-tag was then cleaved off using TEV protease and assessed via SDS-PAGE. C, the AviTag was then biotinylated and RNF4N-biotin purified via gel filtration. D, LCMS analysis was used at each stage of sample production (His[6]-tag cleavage, biotinylation, FRET fluorophore conjugation).

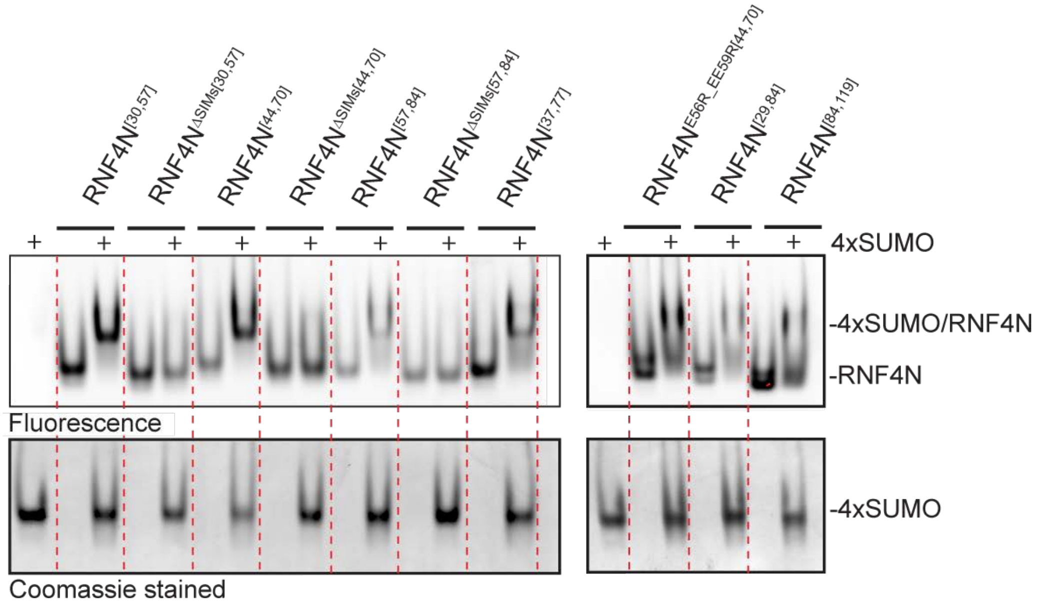

**Extended data figure 2. Binding between RNF4N peptides and 4xSUMO.**

Top panel, in-gel fluorescence analysis of the various RNF4N single-molecule peptides. RNF4N was incubated with/without 4xSUMO and then resolved via native PAGE electrophoresis. Samples incubated with 4xSUMO are highlighted above the gel image. Following in-gel fluorescence analysis the gels were then Coomassie stained to highlight the 4xSUMO, bottom panel.

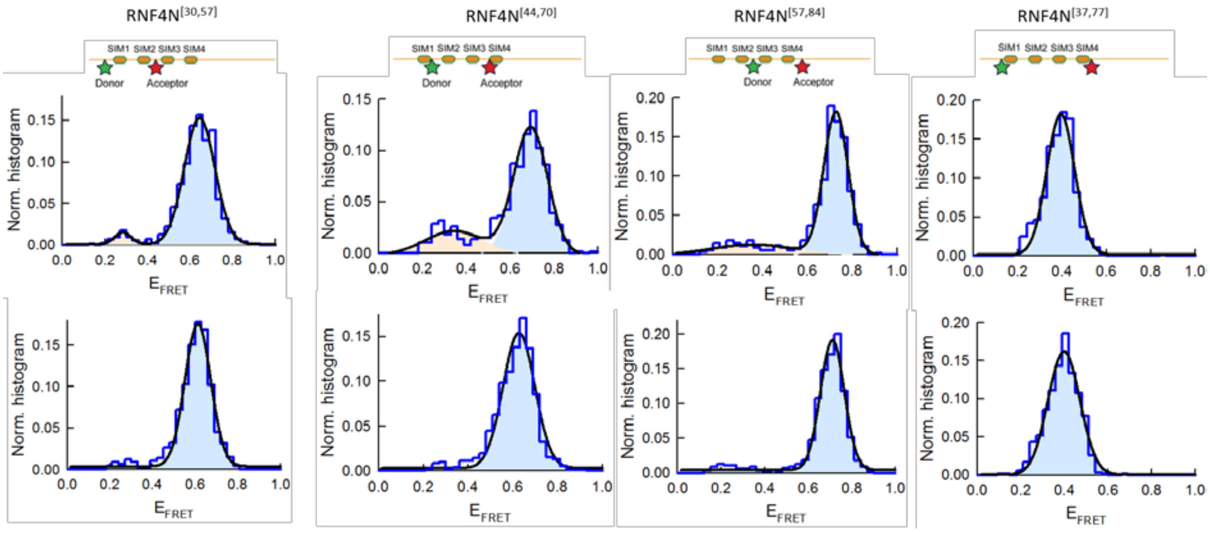

**Extended data figure 3. Single-molecule FRET histograms with Gaussian fit.**

Single-molecule FRET histograms (dark blue) for the SIM peptides carrying the donor (green) and acceptor (red) at the specified positions in the absence (top panels) and in the presence (bottom panels) of SUMO. The histograms are similar to those shown in Figure 1 C but they have been fitted to one or two Gaussians depending on the specific peptide and representing the distribution of FRET populations. Each Gaussian is represented by orange and light blue coloured areas. The solid black line represents the sum of gaussian populations. Single-molecule FRET histograms have been normalized to area unity.

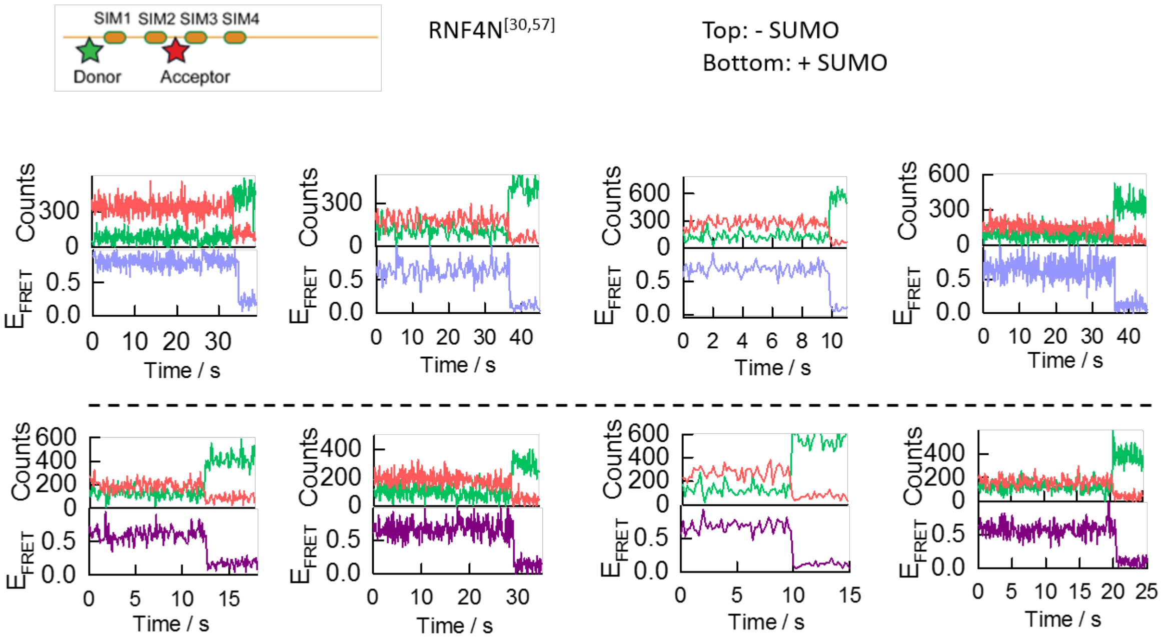

**Extended data figure 4. Representative single-molecule trajectories.**

Representative single-molecule donor (green) and acceptor (red) intensity trajectories (top panels) and corresponding single-molecule FRET trace (bottom panels) obtained for the RNF4 30/57 peptide in the presence (light blue) and absence (purple) of SUMO.

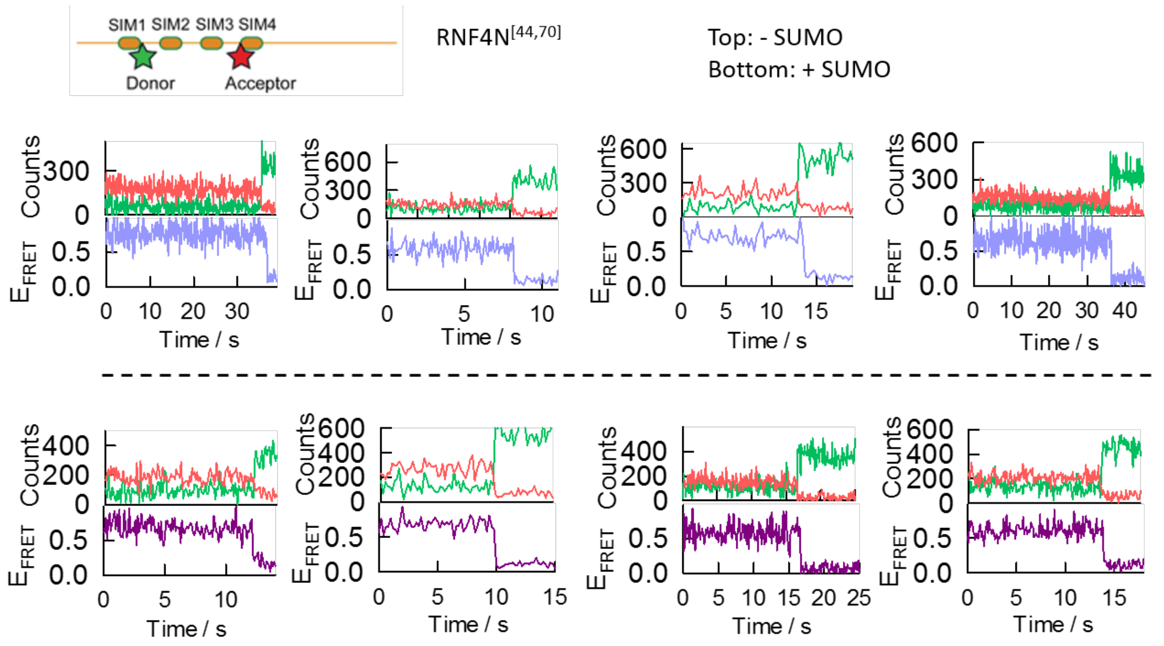

**Extended data figure 5. Representative single-molecule trajectories.**

Representative single-molecule donor (green) and acceptor (red) intensity trajectories (top panels) and corresponding single-molecule FRET trace (bottom panels) obtained for the RNF4 40/70 peptide in the presence (light blue) and absence (purple) of SUMO.

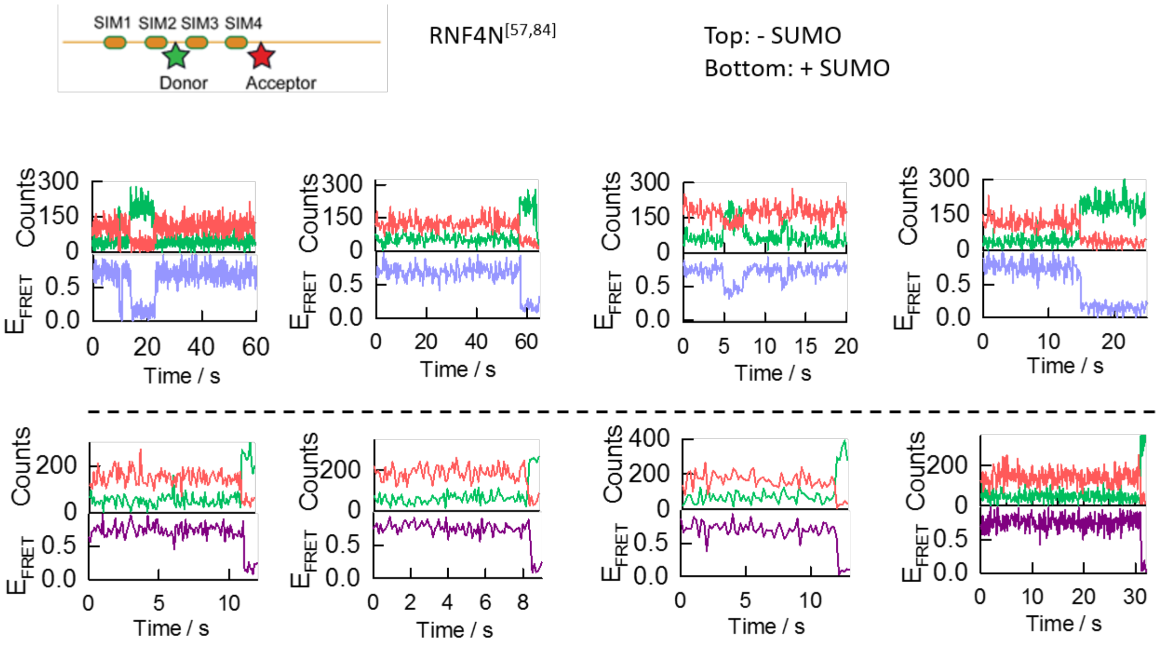

**Extended data figure 6. Representative single-molecule trajectories.**

Representative single-molecule donor (green) and acceptor (red) intensity trajectories (top panels) and corresponding single-molecule FRET trace (bottom panels) obtained for the RNF4 57/84 peptide in the presence (light blue) and absence (purple) of SUMO.

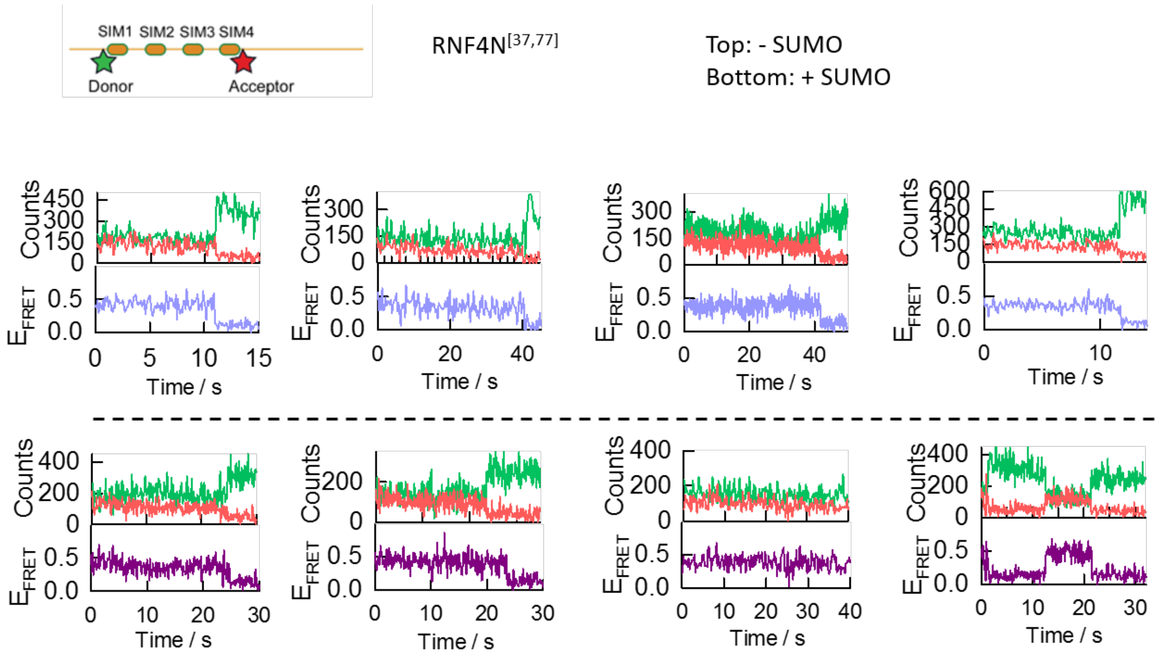

**Extended data figure 7. Representative single-molecule trajectories.**

Representative single-molecule donor (green) and acceptor (red) intensity trajectories (top panels) and corresponding single-molecule FRET trace (bottom panels) obtained for the RNF4 37/77 peptide in the presence (light blue) and absence (purple) of SUMO.

**Table 1. Analysis of histograms Extended data figure 3.**

FRET efficiencies, relative populations and distribution-width values obtained with and without SUMO for RNF4N 30/57, RNF4N 44/70, RNF4N 57/84 and RNF4N 37/77 peptides. Values were calculated by non-linear squares fitting of the experimental single-molecule FRET histograms shown in Figure 1C and Figure ED3 to one or two Gaussian distributions describing the number and relative contribution of FRET populations. The $E_{FRET}$ magnitude was taken from the centre of the Gaussian and the error represents the standard error of the fit. The width of each Gaussian and its relative contribution are also shown. The value of $\Delta E_{FRET}$ was calculated only for the most populated FRET state as the difference between $E_{FRET}$ in the presence of SUMO and $E_{FRET}$ with no SUMO added. Colours match those shown in Figure ED3 for each FRET population.

| SIM peptides | $E_{FRET}$ | Width | % | $\Delta E_{FRET}$ | $E_{FRET}$ | Width | % |
|---|---|---|---|---|---|---|---|
| **RNF4N 30/57** | 0.642 ± 0.002 | 0.141 ± 0.002 | 95.2 | -0.029 ± 0.003 | 0.28 ± 0.02 | 0.08± 0.02 | 4.8 |
| **RNF4N 30/57 + SUMO** | 0.613 ± 0.002 | 0.123 ± 0.032 | 100 | | -- | -- | -- |
| **RNF4N 44/70** | 0.693 ± 0.004 | 0.153 ± 0.001 | 75.5 | -0.066 ± 0.005 | 0.34 ± 0.03 | 0.26± 0.07 | 24.5 |
| **RNF4N 44/70 + SUMO** | 0.627 ± 0.003 | 0.141 ± 0.006 | 100 | | -- | -- | -- |
| **RNF4N 57/84** | 0.727 ± 0.002 | 0.108 ± 0.004 | 81 | -0.016 ± 0.003 | 0.35 ± 0.05 | 0.36 ± 0.15 | 19 |
| **RNF4N 57/84 + SUMO** | 0.711 ± 0.002 | 0.107 ± 0.004 | 100 | | -- | -- | -- |
| **RNF4N 37/77** | 0.394 ± 0.002 | 0.126 ± 0.005 | 100 | 0.004 ± 0.003 | -- | -- | -- |
| **RNF4N 37/77 + SUMO** | 0.398 ± 0.002 | 0.146 ± 0.005 | 100 | | -- | -- | -- |

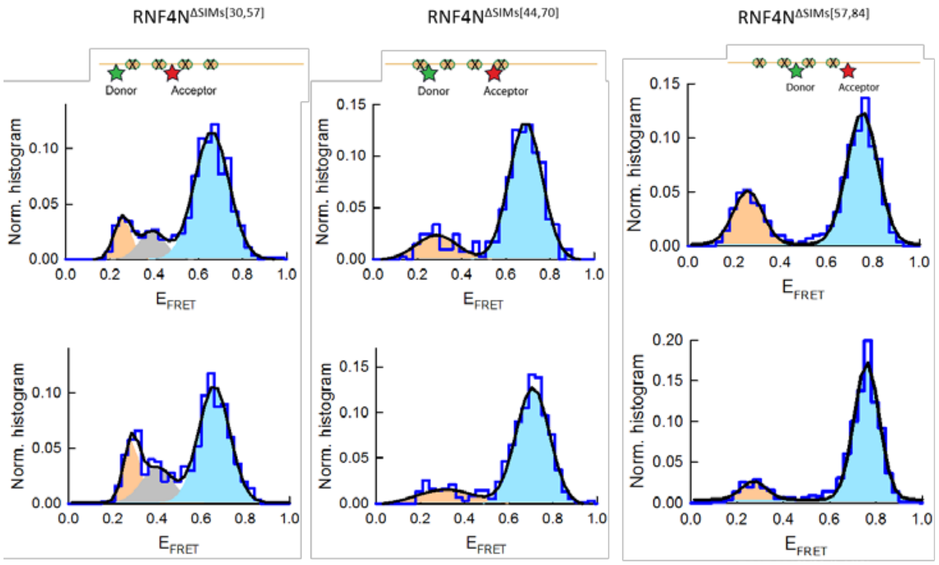

**Extended data figure 8. Single-molecule FRET histograms with Gaussian fit.**

Single-molecule FRET histograms (dark blue) for the SIM mutant peptides carrying the donor (green) and acceptor (red) at the specified positions in the absence (top panels) and in the presence (bottom panels) of SUMO. The histograms are similar to those shown in Figure 2 A but they have been fitted to one, two or three Gaussians depending on the specific peptide and representing the distribution of FRET populations. The contribution of each FRET population is represented by orange, grey and light blue coloured areas. The solid black line represents the sum of Gaussian populations. Single-molecule FRET histograms have been normalized to area unity.

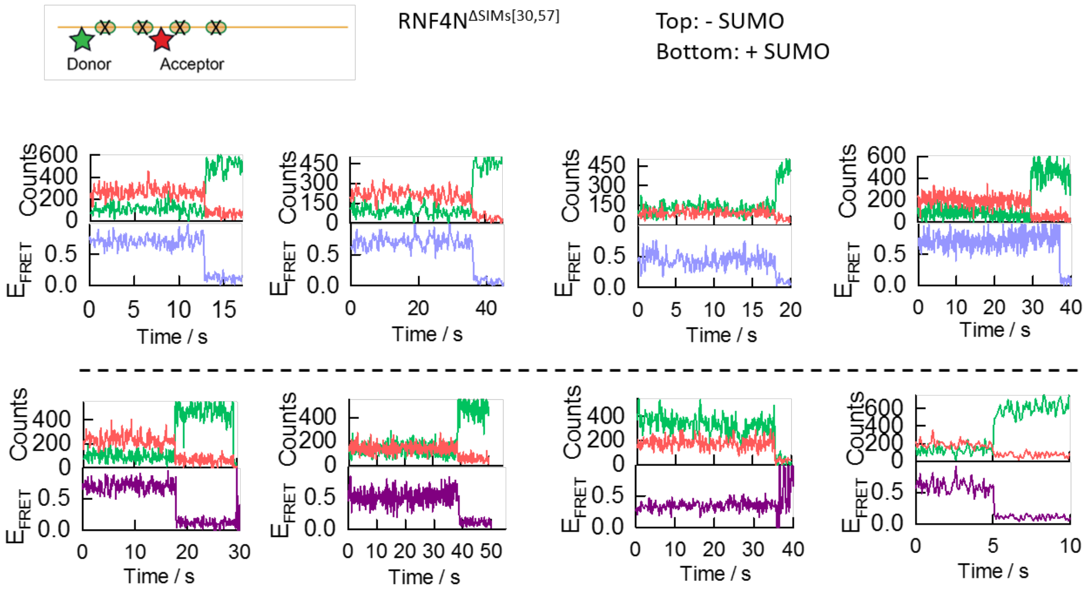

**Extended data figure 9. Representative single-molecule trajectories.**

Representative single-molecule donor (green) and acceptor (red) intensity trajectories (top panels) and corresponding single-molecule FRET trace (bottom panels) obtained for the RNF4N$^{\Delta SIMs}$ 30/57 peptide in the presence (light blue) and absence (purple) of SUMO.

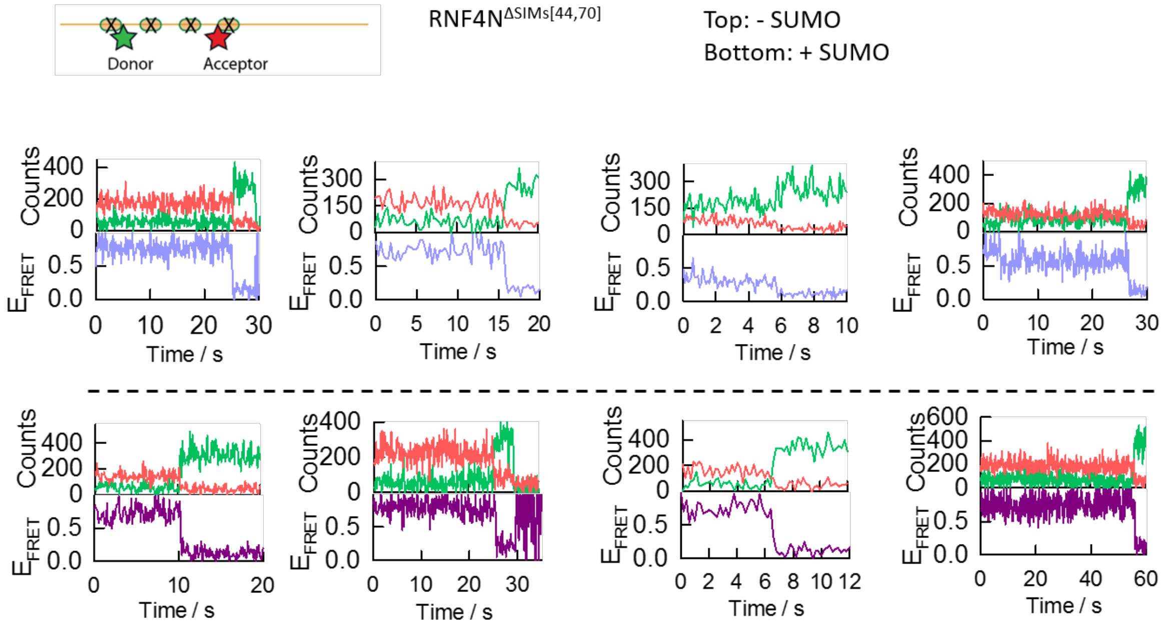

**Extended data figure 10. Representative single-molecule trajectories.**

Representative single-molecule donor (green) and acceptor (red) intensity trajectories (top panels) and corresponding single-molecule FRET trace (bottom panels) obtained for the RNF4N$^{\Delta SIMs}$ 44/70 peptide in the presence (light blue) and absence (purple) of SUMO.

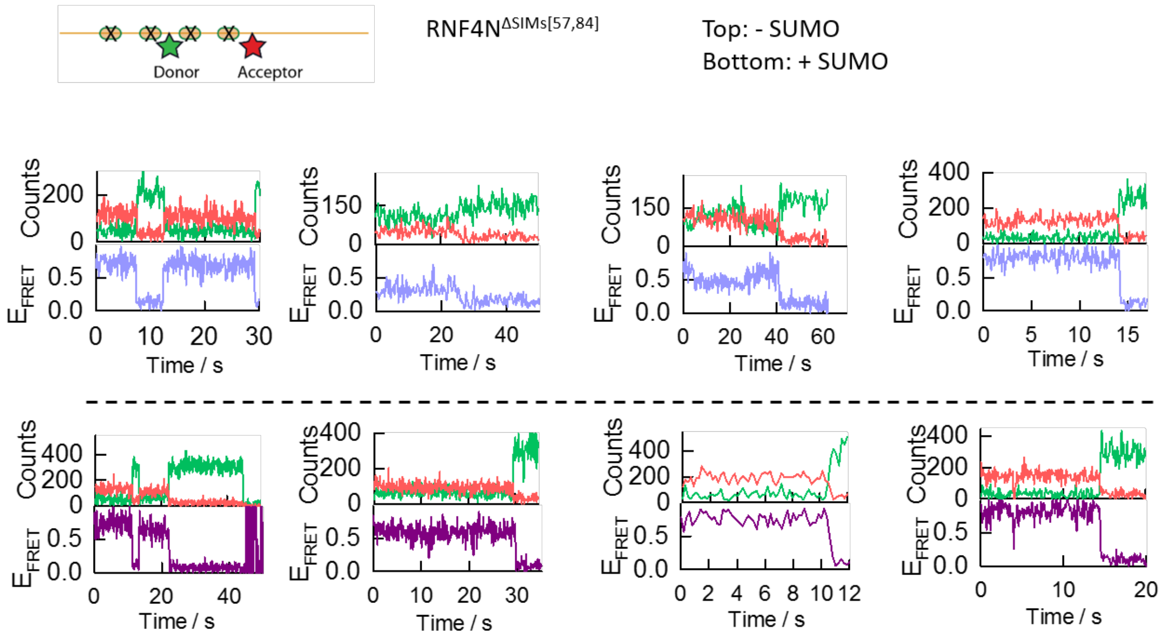

**Extended data figure 11. Representative single-molecule trajectories.**

Representative single-molecule donor (green) and acceptor (red) intensity trajectories (top panels) and corresponding single-molecule FRET trace (bottom panels) obtained for the RNF4N$^{\Delta SIMs}$ 57/84 peptide in the presence (light blue) and absence (purple) of SUMO.

**Table 2. Analysis of histograms Extended data figure 8.**

FRET efficiencies, relative populations and distribution-width values obtained with and without SUMO for SIM mutant peptides RNF4N$^{\Delta SIMs}$ 30/57, RNF4N$^{\Delta SIMs}$ 44/70 and RNF4N$^{\Delta SIMs}$ 57/84. Values were calculated by non-linear squares fitting of the experimental single-molecule FRET histograms shown in Figure 2 A and Figure ED8 to one, two or three Gaussian distributions describing the number and relative contribution of FRET populations. The $E_{FRET}$ magnitude was taken from the centre of the Gaussian and the error represents the standard error of the fit. The width of each Gaussian and its relative contribution are also shown. The value of $\Delta E_{FRET}$ was calculated only for the most populated FRET state as the difference between $E_{FRET}$ in the presence of SUMO and $E_{FRET}$ with no SUMO added. Colours match those shown in Figure ED8 for each FRET population.

| Mutant peptides | $E_{FRET}$ | Width | % | $\Delta E_{FRET}$ | $E_{FRET}$ | Width | %$^2$ | $E_{FRET}$ | Width | % |
|---|---|---|---|---|---|---|---|---|---|---|
| **RNF4N$^{\Delta SIMs}$ 30/57** | 0.661 ± 0.003 | 0.157 ± 0.008 | 75.4 | -0.002 ± 0.005 | 0.26 ± 0.01 | 0.08 ± 0.02 | 12.1 | 0.39 ± 0.02 | 0.12 ± 0.05 | 12.5 |
| **RNF4N$^{\Delta SIMs}$ 30/57 + SUMO** | 0.659 ± 0.004 | 0.14 ± 0.01 | 64.1 | | 0.28 ± 0.01 | 0.07 ± 0.01 | 16.9 | 0.40 ± 0.04 | 0.14 ± 0.08 | 19 |
| **RNF4N$^{\Delta SIMs}$ 44/70** | 0.689 ± 0.003 | 0.149 ± 0.007 | 81.7 | +0.02 ± 0.004 | 0.29 ± 0.02 | 0.18 ± 0.05 | 18.3 | -- | -- | -- |
| **RNF4N$^{\Delta SIMs}$ 44/70 + SUMO** | 0.709 ± 0.003 | 0.152 ± 0.006 | 81.1 | | 0.32 ± 0.03 | 0.27 ± 0.07 | 18.9 | -- | -- | -- |
| **RNF4N$^{\Delta SIMs}$ 57/84** | 0.754 ± 0.002 | 0.133 ± 0.005 | 72.3 | -0.005 ± 0.003 | 0.259 ± 0.006 | 0.12 ± 0.01 | 27.7 | -- | -- | -- |
| **RNF4N$^{\Delta SIMs}$ 57/84 + SUMO** | 0.759 ± 0.002 | 0.108 ± 0.005 | 88.1 | | 0.27 ± 0.02 | 0.11 ± 0.04 | 11.9 | -- | -- | -- |

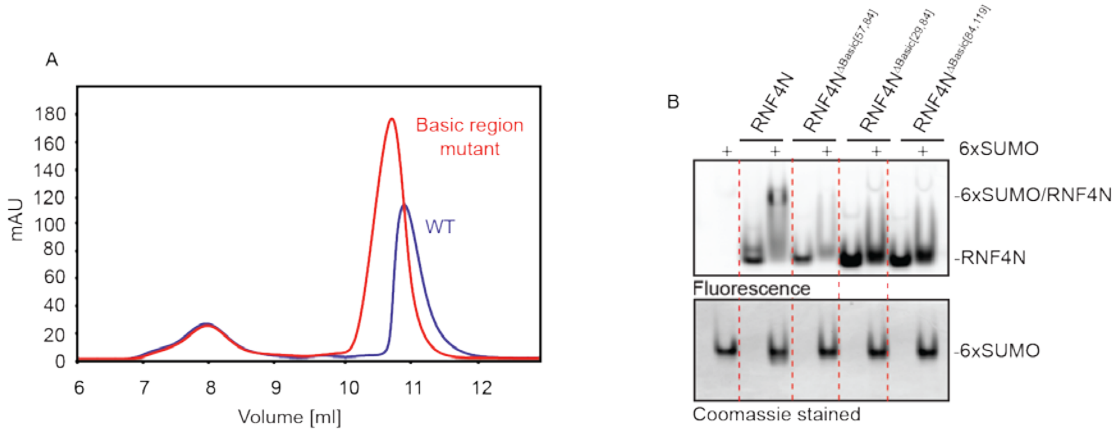

**Extended data figure 12. Assessment of basic region mutations to RNF4N peptides.**

**A,** gel filtration of RNF4N peptide with removal of basic charge from the SIMs-RING linker region causes early elution compared with wild type RNF4N. **B**, top panel in-gel fluorescence analysis of the various RNF4N basic region mutant single-molecule peptides. RNF4N was incubated with/without 6xSUMO and then resolved via native PAGE electrophoresis. Samples incubated with 6xSUMO are highlighted above the gel image. Following in-gel fluorescence analysis the gels were then Coomassie stained to highlight the 6xSUMO, bottom panel.

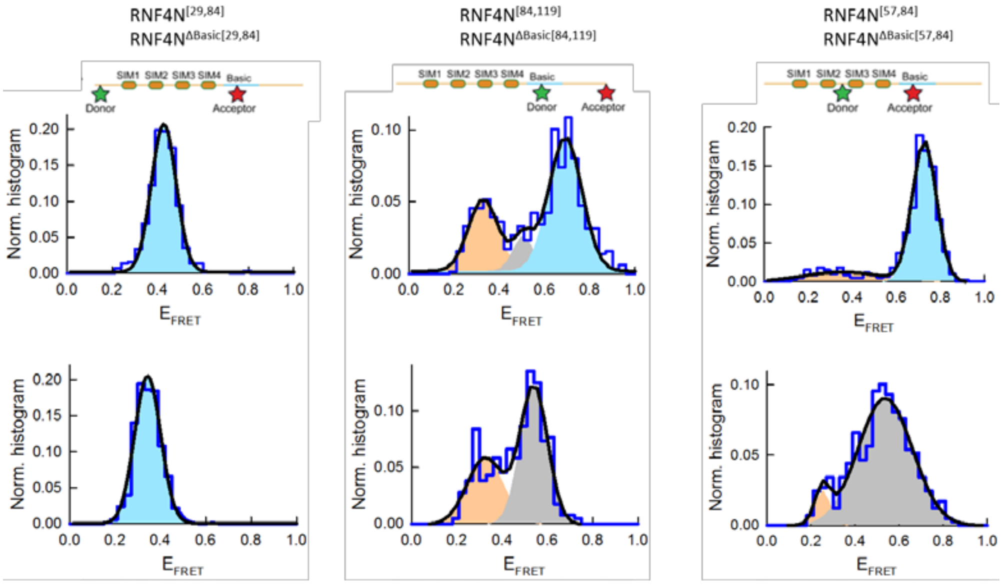

**Extended data figure 13. Single-molecule FRET histograms with Gaussian fit.**

Single-molecule FRET histograms (dark blue) for the SIM mutant peptides carrying the donor (green) and acceptor (red) at the specified positions. The histograms are similar to those shown in Figure 2 C but they have been fitted to one, two or three gaussians depending on the specific peptide and representing the distribution of FRET populations. The contribution of each FRET population is represented by orange, grey and light blue coloured areas. The solid black line represents the sum of gaussian populations. Single-molecule FRET histograms have been normalized to area unity.

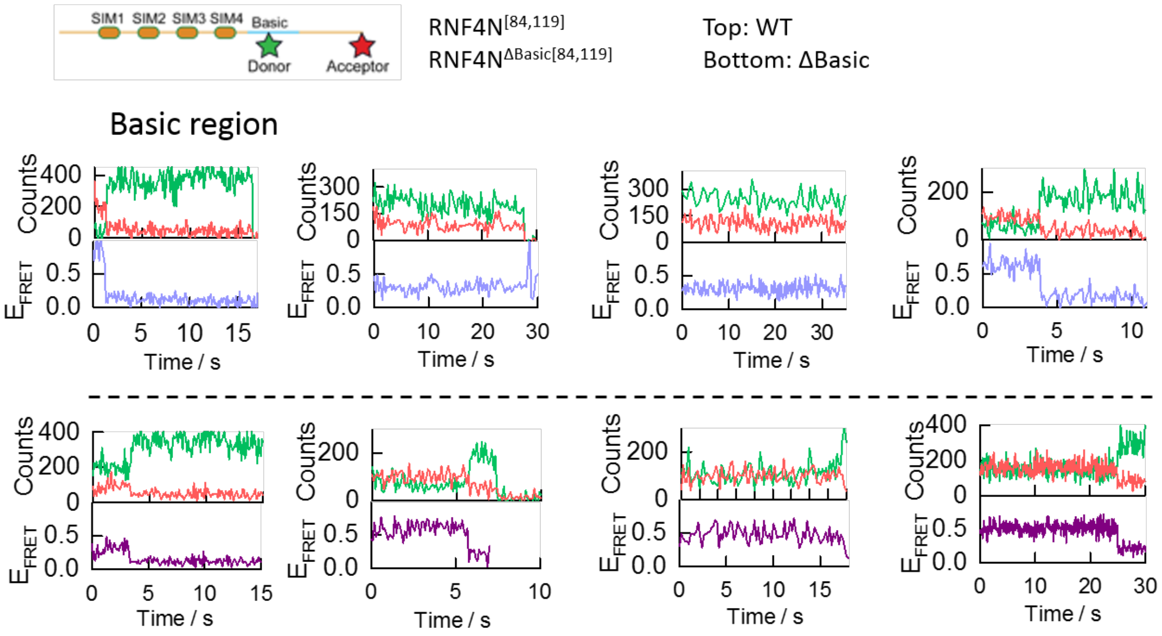

**Extended data figure 14. Representative single-molecule trajectories.**

Representative single-molecule donor (green) and acceptor (red) intensity trajectories (top panels) and corresponding single-molecule FRET trace (bottom panels) obtained for the RNF4N$^{\Delta basic}$ 84/119 peptide (light blue) and mutated version (purple).

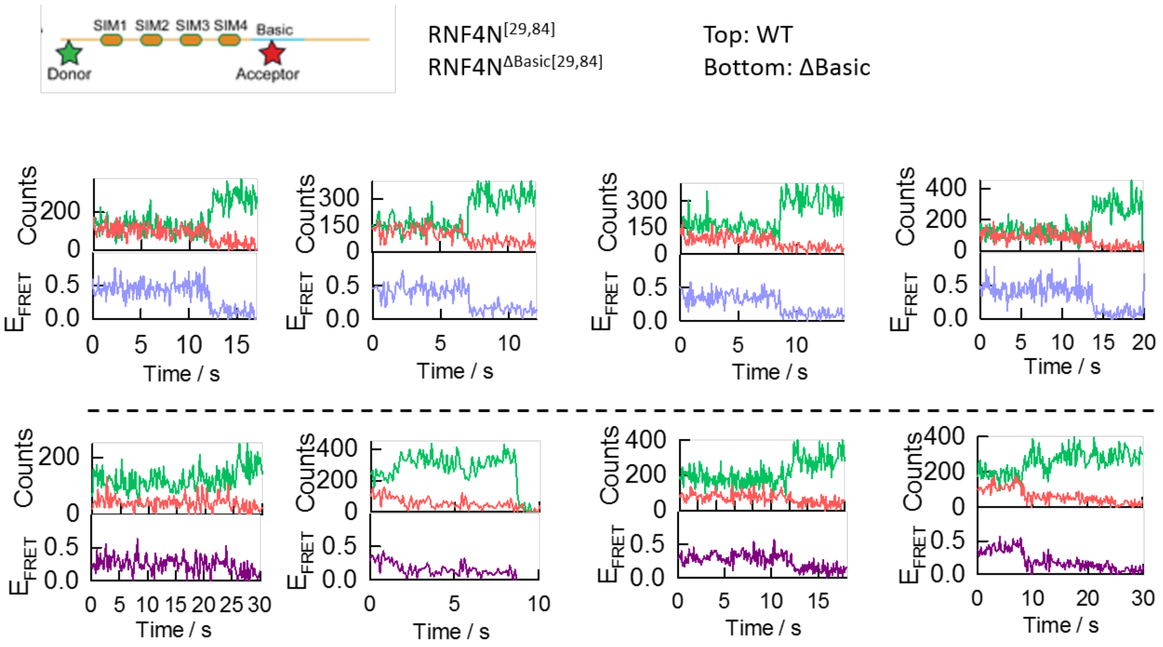

**Extended data figure 15. Representative single-molecule trajectories.**

Representative single-molecule donor (green) and acceptor (red) intensity trajectories (top panels) and corresponding single-molecule FRET trace (bottom panels) obtained for the RNF4N$^{\Delta basic}$ 29/84 peptide (light blue) and mutated version (purple).

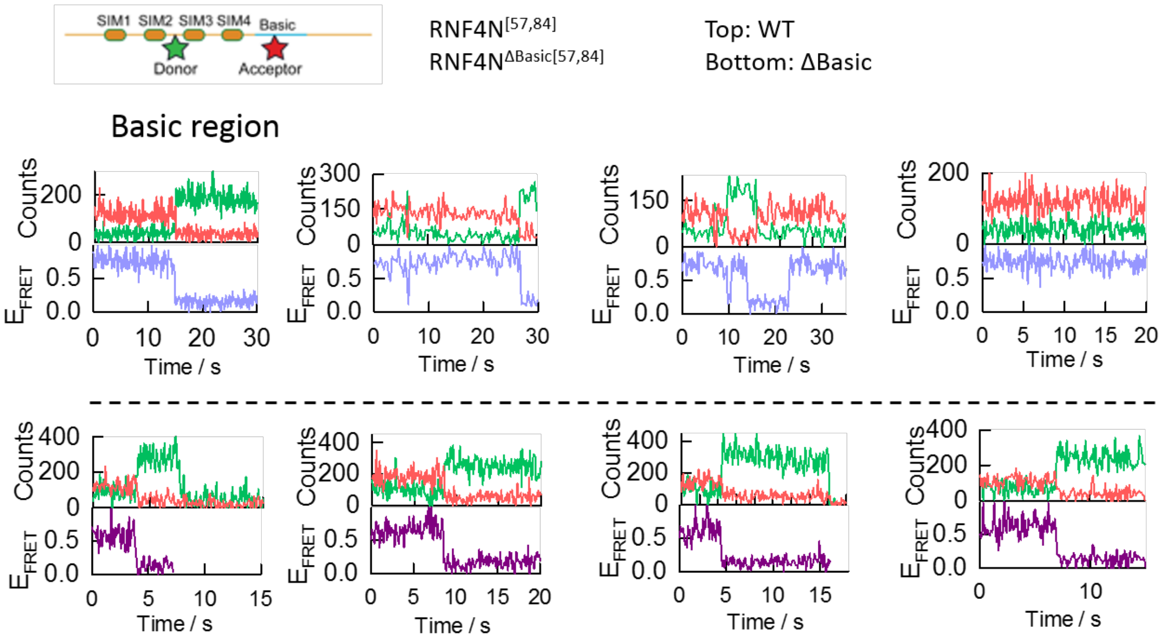

**Extended data figure 16. Representative single-molecule trajectories.**

Representative single-molecule donor (green) and acceptor (red) intensity trajectories (top panels) and corresponding single-molecule FRET trace (bottom panels) obtained for the RNF4N$^{\Delta basic}$ 57/84 peptide (light blue) and mutated version (purple).

**Table 3. Analysis of histograms Extended data figure 13.**

FRET efficiencies, relative populations and distribution-width values obtained for basic region mutant peptides RNF4$^{\Delta Basic}$ 29/84, RNF4$^{\Delta Basic}$ 84/119 and RNF4N$^{\Delta Basic}$ 57/84. Values were calculated by non-linear squares fitting of the experimental single-molecule FRET histograms shown in Figure 2C and Figure ED13 to one, two or three Gaussian distributions describing the number and relative contribution of FRET populations. The $E_{FRET}$ magnitude was taken from the centre of the Gaussian and the error represents the standard error of the fit. The width of each Gaussian and its relative contribution are also shown. The value of $\Delta E_{FRET}$ was calculated as the difference between the $E_{FRET}$ for the most populated FRET state at each experimental condition. Colours match those shown in Figure ED13 for each FRET population.

| Mutant peptides | $E_{FRET}$ | Width | % | $\Delta E_{FRET}$ | $E_{FRET}$ | Width | % | $E_{FRET}$ | Width | % |
|---|---|---|---|---|---|---|---|---|---|---|
| **RNF4 29/84** | 0.427 ± 0.001 | 0.109 ± 0.002 | 100 | -0.084 ± 0.002 | -- | -- | -- | -- | -- | -- |
| **RNF4$^{\Delta Basic}$ 29/84** | 0.343 ± 0.002 | 0.115 ± 0.003 | 100 | | | | | | | |
| **RNF4 84/119** | 0.691 ± 0.001 | 0.15 ± 0.01 | 60.9 | -0.15 ± 0.006 | 0.33 ± 0.01 | 0.13 ± 0.03 | 29.4 | 0.50 ± 0.02 | 0.09 ± 0.04 | 9.6 |
| **RNF4$^{\Delta Basic}$ 84/119** | 0.32 ± 0.01 | 0.16 ± 0.03 | 38.9 | | | | | 0.540 ± 0.006 | 0.12 ± 0.01 | 61.1 |
| **RNF4N 57/84** | 0.727 ± 0.002 | 0.108 ± 0.004 | 81 | -0.19 ± 0.006 | 0.35 ± 0.05 | 0.36 ± 0.15 | 19 | | | |
| **RNF4$^{\Delta Basic}$ 57/84** | | | | | 0.25 ± 0.01 | 0.07 ± 0.02 | 7 | 0.537 ± 0.006 | 0.24 ± 0.01 | 93 |

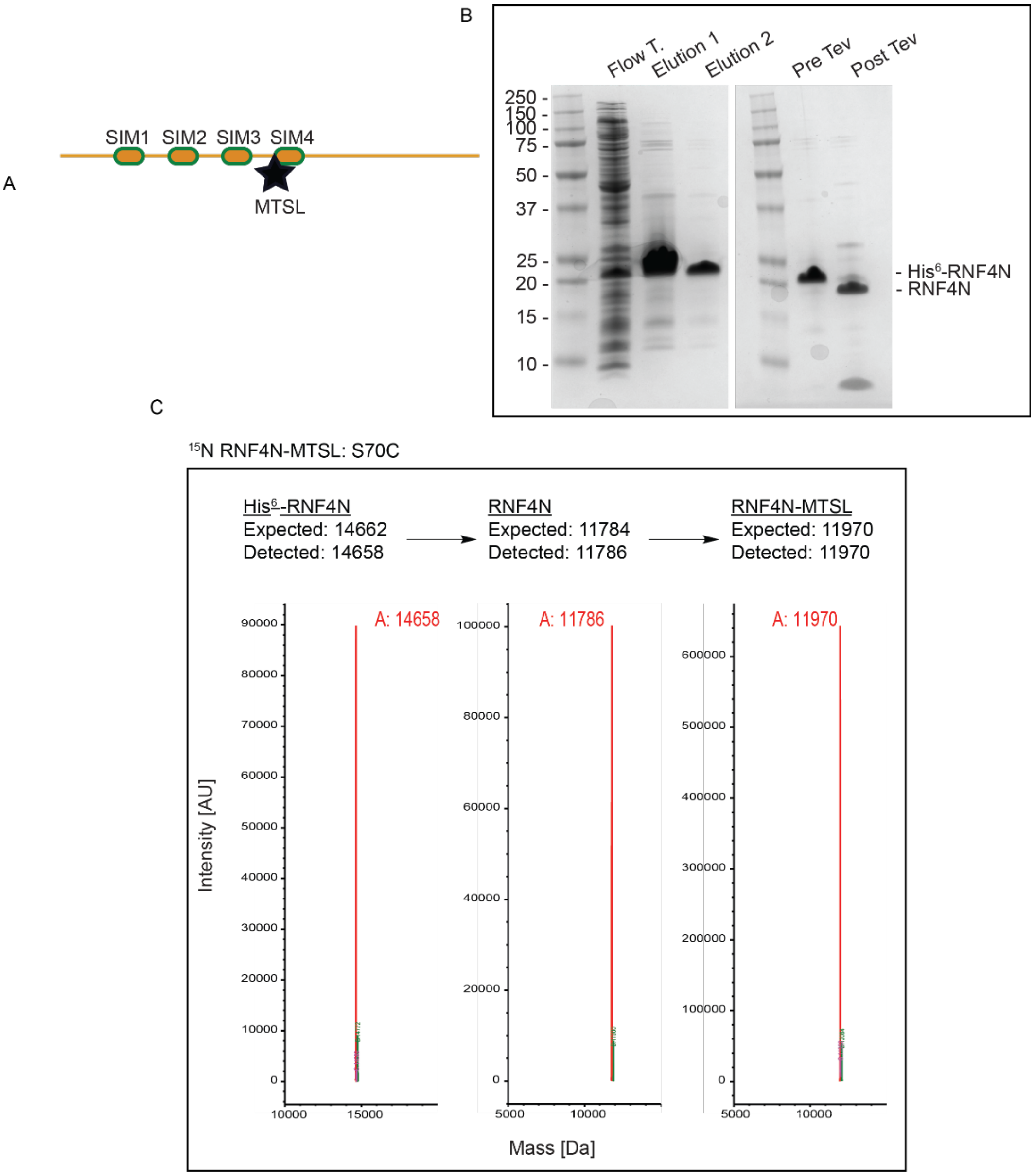

**Extended data figure 17. Production of $^{15}$N-enriched RNF4N with MTSL spin label.**

A, model of the N-terminal region of RNF4 (RNF4N) bearing a MTSL spin label next to SIM4. B, RNF4 was initially expressed with an N-terminal His$^6$-tag to allow for purification. The His$^6$-tag is was then cleaved off using TEV protease and assessed via SDS-PAGE. C, LCMS analysis was used at each stage of sample production (His$^6$-tag cleavage, MTSL labelling).

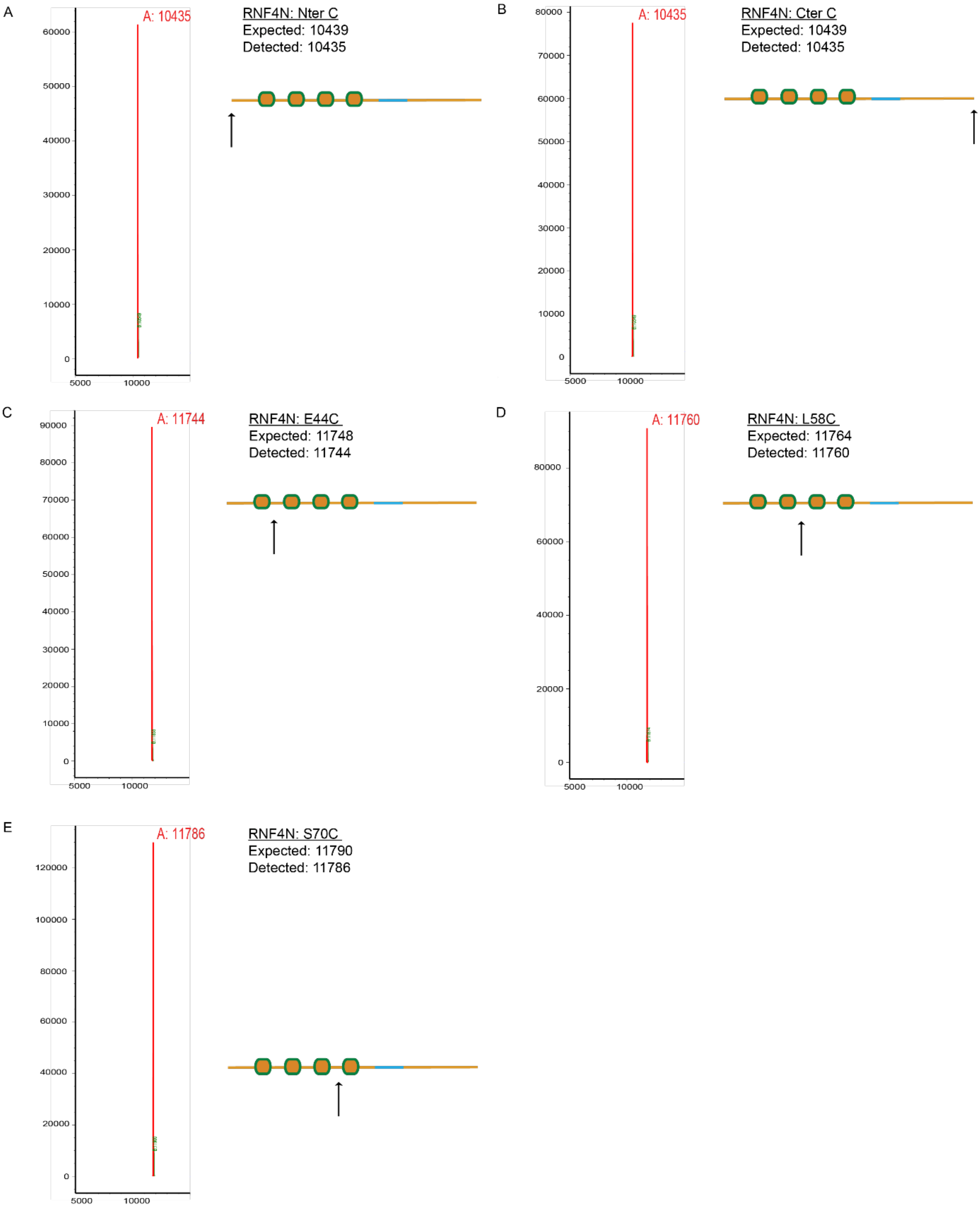

**Extended data figure 18. LCMS validation of all $^{15}$N-enriched RNF4N peptides labelled with MTSL**

Five different 15N-enriched RNF4N peptides were produced, each containing on cysteine residue for site-directed attachment of an MTSL spin label. Following spin-label addition the mass of each peptide was validated via LCMS. Illustrations inset highlight the position of spin label attachment for each peptide.

**Table 4. Properties calculated from molecular dynamics simulations from figure 3 D.**

Rg = radius of gyration; % SAS is the percentage change in protein surface area compared to the fully extended conformation, end to end distance is measured between the termini, and middle-to-end distance is calculated between residue 80 and residue 133. All values were calculated as a mean over the final 5 μs of simulation data with the standard deviation shown in parenthesis.

| | Run number | Rg | % change buried surface area | Distance end-to-end | Distance mid-to-end |
|---|---|---|---|---|---|
| Wildtype | 1 | 6.6 (0.5) | 3.1 (0.6) | 7.2 (2.6) | 10.7 (1.7) |
| | 2 | 2.9 (0.3) | 9.5 (1.1) | 4.6 (2.1) | 5.4 (1.5) |
| | 3 | 3.6 (0.8) | 8.6 (4.3) | 9.6 (1.3) | 4.2 (0.9) |
| | 4 | 4.5 (0.5) | 5.1 (5.4) | 12.9 (2.4) | 6.5 (0.8) |
| | 5 | 3.7 (0.4) | 6.1 (5.0) | 8.1 (1.7) | 7.6 (0.9) |
| | overall | 4.1 (1.4) | 6.3 (2.8) | 8.5 (3.4) | 6.9 (2.5) |
| Basic mutant | 1 | 5.1 (0.5) | 3.8 (1.2) | 9.1 (1.8) | 8.6 (2.2) |
| | 2 | 4.3 (0.8) | 12.8 (2.3) | 9.4 (1.9) | 6.2 (0.4) |
| | 3 | 6.4 (0.7) | 3.0 (1.0) | 14.9 (2.3) | 10.2 (2.0) |
| | 4 | 5.9 (0.5) | 2.9 (1.0) | 9.5 (3.2) | 10.1 (1.6) |
| | 5 | 3.2 (0.4) | 12.5 (7.8) | 7.3 (1.8) | 6.0 (1.2) |
| | overall | 5.0 (1.3) | 7.0 (3.9) | 10.1 (3.4) | 8.3 (2.4) |

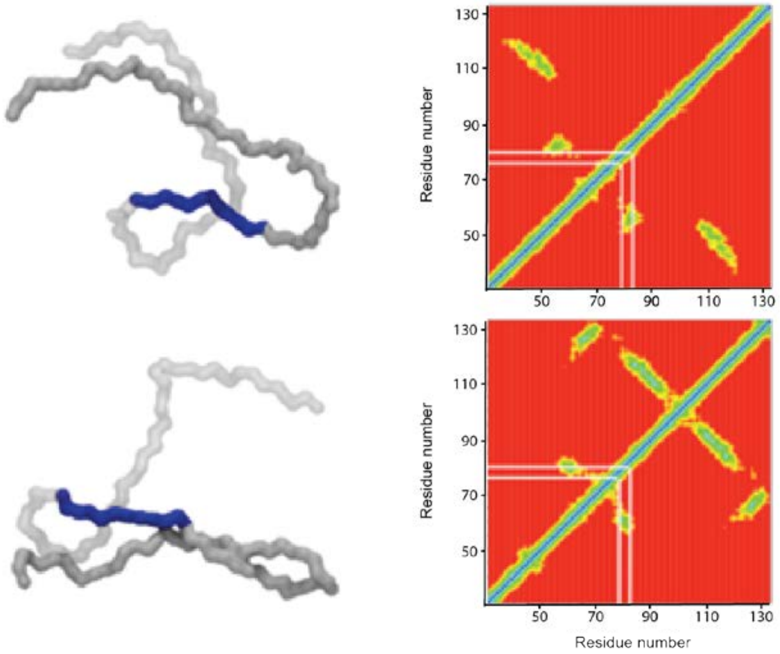

**Extended data figure 19. Contact plots displaying electrostatic interactions.**

As shown in figure 3 D, the contact plots show the residue-residue (X-Y axis) contacts for wild type RNF4 peptide (residues 32-133). Electrostatic interactions involving the charged basic region is highlighted by white lines. These contacts are not observed in the mutant simulations.

The peptide is coloured: basic region residues 77-86 (blue), N-terminal residues 32-76 (light grey), C-terminal residues 87-133 (dark grey).

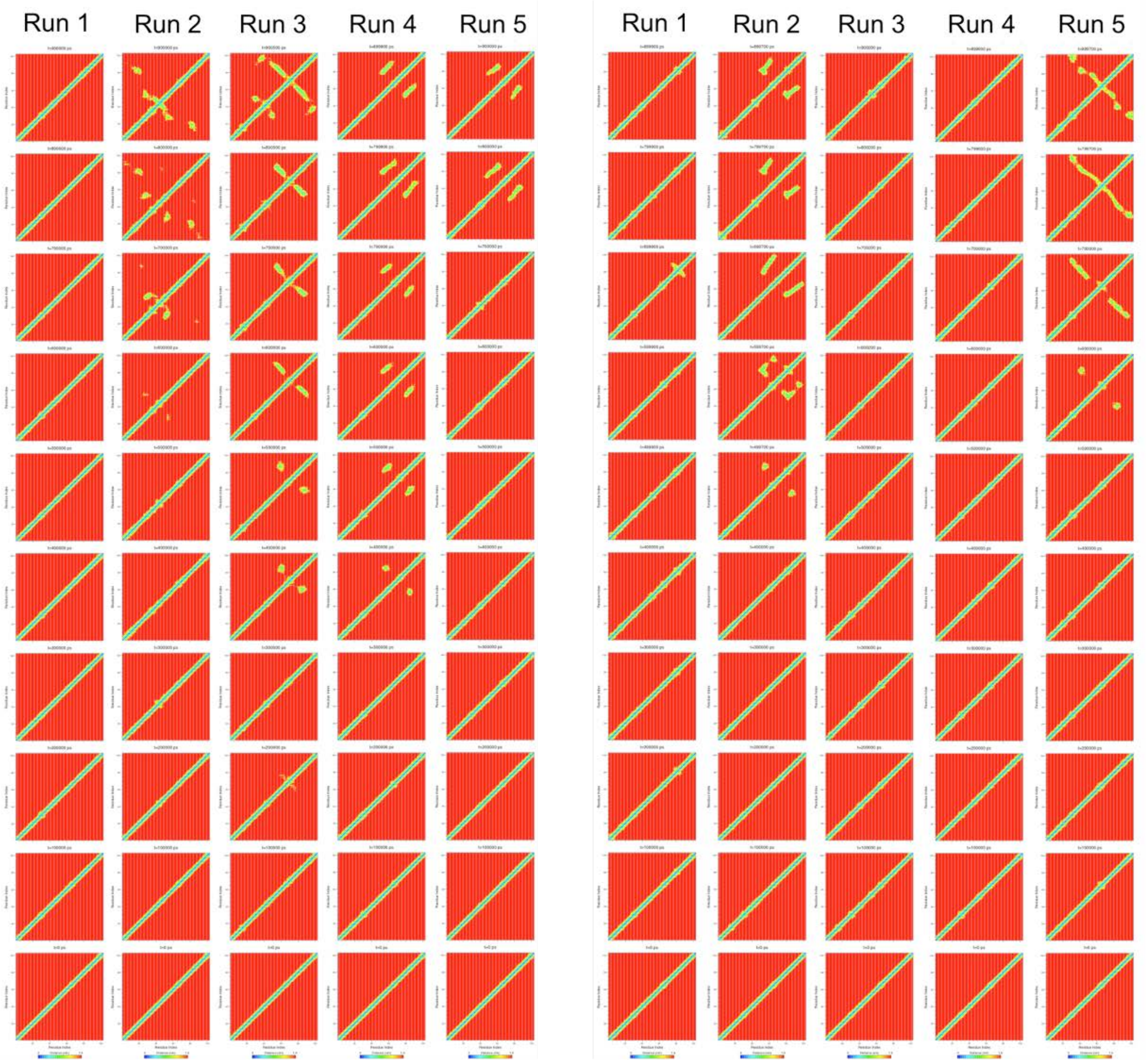

**Extended data figure 19**. **Contact plots.**

Extended contact plots of RNF4 wild type (left hand series) and basic mutant (right hand series) from simulation trajectories, as seen in figure 3 D. Protein residue-residue contacts were calculated at 1 microsecond intervals over each trajectory. The contact plots show the residue-residue (X-Y axis) contacts for residues 32-133 of RNF4. Time course starts at the bottom, with the top panel in each run representing the end of the simulation.

**Table 5. Description of RNF4 used in different experimental analysis.**

| Construct | Analysis type | Frame | Description |
|---|---|---|---|
| RNF4N | Single-molecule FRET | residues 27-118 | N-terminal region of RNF4. Contains a C-terminal AviTag |
| RNF4N | NMR | residues 30-118 or 32-133 | N-terminal region of RNF4. Peptides labelled at E44, L58, S70 are 32-133 |
| RNF4 | biochemical analysis | Full length | C55S and C95S mutations are present in WT and basic region mutant |

**Table 6. Description of RNF4N variants including mutations and attachment positon of function modifications (FRET dyes, spin labels).**

| SM peptides | Labelled res. | Labelling description | Additional modifications |
|---|---|---|---|
| RNF4N 30/57 | T30C, S57C | FRET dyes span SIM1+2 | |
| RNF4N$^{\Delta SIMs}$ 30/57 | T30C, S57C | FRET dyes span SIM1+2 | All SIMs mutated |
| RNF4N 44/70 | E44C, S70C | FRET dyes span SIM2+3 | |
| RNF4N$^{\Delta SIMs}$ 44/70 | E44C, S70C | FRET dyes span SIM2+3 | All SIMs mutated |
| RNF4N 57/84 | S57C, G84C | FRET dyes span SIM3+4 | |
| RNF4N$^{\Delta SIMs}$ 57/84 | S57C, G84C | FRET dyes span SIM3+4 | All SIMs mutated |
| RNF4N$^{\Delta Basic}$ 57/84 | S57C, G84C | FRET dyes span SIM3+4 | Basic residues mutated to serine/alanine |
| RNF4N 37/77 | A37C ,R77C | FRET dyes span SIM1-4 | |
| RNF4N 29/84 | S29C, G84C | FRET dyes: N-terminus + basic region | |
| RNF4N$^{\Delta Basic}$ 29/84 | S29C, G84C | FRET dyes: N-terminus + basic region | Basic residues mutated to serine/alanine |
| RNF4N 84/119 | G84C, S119C | FRET dyes: C-terminus + basic region | |
| RNF4N$^{\Delta Basic}$ 84/119 | G84C, S119C | FRET dyes: C-terminus + basic region | Basic residues mutated to serine/alanine |
| **NMR peptides** | | | |
| RNF4N 29 | S29C | Spin label at N-terminus | |
| RNF4N 119 | S119C | Spin label at C-terminus | |
| RNF4N 44 | E44C | Spin label between SIM1+2 | |
| RNF4N 58 | L58C | Spin label between SIM2+3 | |
| RNF4N 70 | S70C | Spin label between SIM3+4 | |
